# Oxytocin promotes advantageous inequity preference by modulating the fairness norm enforcement system

**DOI:** 10.64898/2026.08.11.744311

**Authors:** Shuxia Yao, Yanan Qing, Jiayuan Wang, Yuan Zhang, Hongrui Lin, Haochen Zou, Qiong Zhang, Keith M. Kendrick

**Affiliations:** The Center of Psychosomatic Medicine, Sichuan Provincial Center for Mental Health, Sichuan Provincial People’s Hospital, University of Electronic Science and Technology of China, Chengdu, 611731, China; The Clinical Hospital of Chengdu Brain Science Institute, MOE-K Lab for NeuroInformation, Brain-Apparatus Communication Institute, University of Electronic Science and Technology of China, Chengdu, 611731, China; Brain-Computer Interface & Brain-Inspired Intelligence Key Laboratory of Sichuan Province, Chengdu, China

## Abstract

Aversion to inequity is innate in humans and constitutes a major contributor to social tension. Such aversion can be triggered either when one receives more (advantageous inequity-AI) or less (disadvantageous inequity-DI) than others. However, it remains unclear whether oxytocin can modulate how humans respond to inequity. The present pre-registered study combined behavioral measures, computational modelling, and fMRI in a novel monetary allocation evaluation paradigm to examine oxytocin’s modulation of behavioral and neural responses to inequity. Results showed that oxytocin increased perceived fairness and AI allocation choices via enhancing AI preference independent of gain vs. loss contexts, which was further confirmed by model-based results of oxytocin decreasing AI aversion. In parallel at the neural level, oxytocin increased fairness rating-related activity in regions within fairness-related norm enforcement systems involved in norm violation detection, salience processing, and cognitive control. Multivariate-based representational similarity analyses provided further evidence that oxytocin induced distinct representational patterns of inequity in these core regions. These results suggest that oxytocin promoted AI preference possibly by strengthening regulatory control to support the pursuit of economic self-interest, while simultaneously enhancing the salience of AI allocations. Our study provides new insights into behavioral and neural mechanisms through which oxytocin modulates fairness-related behavior.

## Introduction

As noted by JS Adams in his seminal work, “There can be little doubt that inequity results in dissatisfaction” [1], aversion to inequity—or, in other words, a social preference for equity—is innate in humans. Inequity emerges when individuals perceive discordance of outcomes between their own and others’ effort inputs [1, 2] and constitutes one of the most persistent issues causing social tension and posing challenges to societal development within and across countries [3, 4]. Note that when we talk about inequity in daily life, we primarily refer to the situation of disadvantageous inequity (DI) whereby one’s own payoffs are less than others. However, in psychological and economic conceptualizations, inequity also involves the situation of advantageous inequity (AI) whereby one receives more than others [5]. Aversion to AI emerges later than to DI and is exclusively observed in humans and chimpanzees [6, 7].

Given the disadvantageous imbalance of payoffs, DI allocations normally elicit strong negative emotions like envy and anger [7–9] and consequently higher rejection rates [10]. Neuroimaging meta-analysis studies have shown that receiving DI proposals in the ultimatum game (UG) are linked with activity in a range of brain regions including the anterior insula (aINS), dorsal anterior cingulate cortex (dACC), dorsolateral prefrontal cortex (dlPFC), and ventrolateral PFC (vlPFC) involved in negative emotion processing, conflict monitoring, and cognitive control [11–13]. Rejecting relative to accepting DI allocations has been shown to induce heightened activity similarly in the aINS, dlPFC, dACC and additionally in the lentiform nucleus and putamen [11, 12]. Behavioral and brain responses to DI offers in the UG are also modulated by gain vs. loss contexts such that loss offers are perceived as more unfair and induce a higher rejection rate associated with stronger activity in the negative emotion processing, conflict monitoring, and cognitive control networks [14, 15].

AI proposals by contrast induce more complex emotional reactions [16–18], which are characterized by initial egoism-based pleasure driven by receiving more than others and subsequently a feeling of guilt induced by norm violation [19, 20]. Within a social comparison framework, the egoism-based pleasure may also manifest in a manner of schadenfreude [21–23], which develops as early as 24 months as a response to inequity aversion [24] and predicts developmental changes in equity-related decisions [25]. Neuroimaging studies have shown that AI allocations following a point estimation task induced higher pleasantness associated with stronger ventral striatum (VS) activity than DI allocations, while DI allocations induced increased activity in the dlPFC compared with AI ones [17]. However, AI and DI allocations could induce similar brain response patterns encompassing the aINS, dlPFC, lateral orbitofrontal cortex (lOFC), and putamen when participants could alter their partner’s payoff at personal cost, although they were more satisfied with AI than DI allocations [26]. Similar brain responses in the emotional and cognitive control networks have also been reported in a modified dictator game (DG) when participants serve as a proposer [27], which can be modulated by a feeling of guilt induced prior to the task [28]. For gain vs. loss contexts, participants are more satisfied with gain than loss offers paralleled with enhanced activation in the dlPFC, putamen, and posterior cingulate cortex (PCC) [26]. Similarly in a modified DG, participants are less willing to increase their partner’s payoff by reducing their AI income in the loss relative to the gain context [29].

As an evolutionarily conserved hypothalamic neuropeptide, oxytocin (OT) plays an important role in modulating human behaviors [30–33]. However, to date, there is limited direct evidence that OT could modulate inequity processing, particularly its neural underpinnings, which is surprising given its well-known role in promoting social bonding, cooperation, and conformity to social norms [31–34]. OT has been found to modulate emotional reactions such as envy and anger [35, 36], and guilt and shame [37], which can be induced by inequity, increase emotional empathy [38–40], and promote punishment to social norm violations [41–43]. However, OT can also promote self-benefiting as opposed to altruistic behavior in conflict or moral dilemma contexts [34, 44], as well as lying for personal gain [45], particularly in males. In terms of neurobiological underpinnings, there is also a well-established overlap between brain substrates encoding inequity and oxytocinergic pathways [46, 47]. These findings together point to the possibility that the oxytocinergic system may be an important candidate for the neuromodulatory mechanism of inequity-related behavior. There is also preliminary evidence for intranasally administered OT selectively promoting adherence of fairness norms in proposers under the high AI but not the DI conditions in a modified DG [48]. However, the majority of previous studies, particularly those in the AI context, have employed paradigms in which participants acted as proposers who determined resource allocations [49]. Notably, there is evidence showing that AI aversion disappears when participants act as recipients compared with as proposers [50]. It still leaves open an important question of whether OT can modulate how humans respond to inequity from the perspective of a recipient, which constitutes the most common situation causing social tension in real world, and its neural underpinnings in terms of both AI and DI contexts. Given the difference of payoff imbalance between AI and DI allocations, although both allocations can be perceived as unfair, subjective preferences for them are different and influence inequity-related decisions [10, 26, 49–51]. Furthermore, previous studies have shown that unequal offers in gain vs. loss contexts can induce different behavioral and neural responses [14, 15, 26, 29]. It also remains unclear whether OT has similar or distinct effects on AI and DI in these two contexts and their underlying neural mechanisms. In addition, while previous studies mainly examined acceptance behavior or satisfaction levels to unequal offers in the UG, these could be confounded by factors such as the strategic consideration [52, 53]. There is still a lack of evidence for the relationship between perceived levels of inequity and subjective preference without such strategic considerations and the role of OT in this.

To address these issues, we developed a novel monetary allocation evaluation (MAE) paradigm in which participants acted as recipients, first rating their perceived fairness of and preference for AI or DI allocations determined by a third party, followed by a choice between the proposed allocation and a fair alternative (see Figure 1). By combining behavioral measurements, computational modelling, and fMRI-based univariate and multivariate analyses in the MAE task, the present pharmacological study allowed us to determine both the behavioral and neural mechanisms via which OT acts to modulate how humans respond to AI vs. DI in gain vs. loss contexts. Although OT is conventionally regarded as a prosocial hormone and has been shown to promote adherence to social norms [33, 41–43], OT-induced increases in punishment of norm violations have typically been observed only when participants’ own benefits were affected or when violators were in-group members [41–43, 48], suggesting a self-serving tendency. Complementing this, OT has been shown to facilitate self-benefiting behavior in conflict, moral dilemma, and unsupervised contexts [34, 44, 45]. Consequently, we hypothesized that OT would increase preference for AI allocations but decrease or have no impact on preference for DI ones, particularly given that in the MAE paradigm participants served as recipients. AI aversion has been found to disappear when participants act as proposers [50], although there is no evidence for OT’s effects on this. At the neural level, we predicted that OT would modulate activity within neural circuitry underlying fairness norm enforcement, including the aINS, dACC, dlPFC, and vlPFC, which support negative emotion processing, conflict monitoring, and cognitive control [17, 26–28, 54–57]. OT could also influence reward-related regions (e.g., the striatum or OFC) that encode the value of allocations [26, 54–56, 58]. These effects may be more pronounced in the loss context, as loss offers have been shown to induce stronger negative responses at both behavioral and neural levels [14, 15, 26, 29].

**Figure 1.**
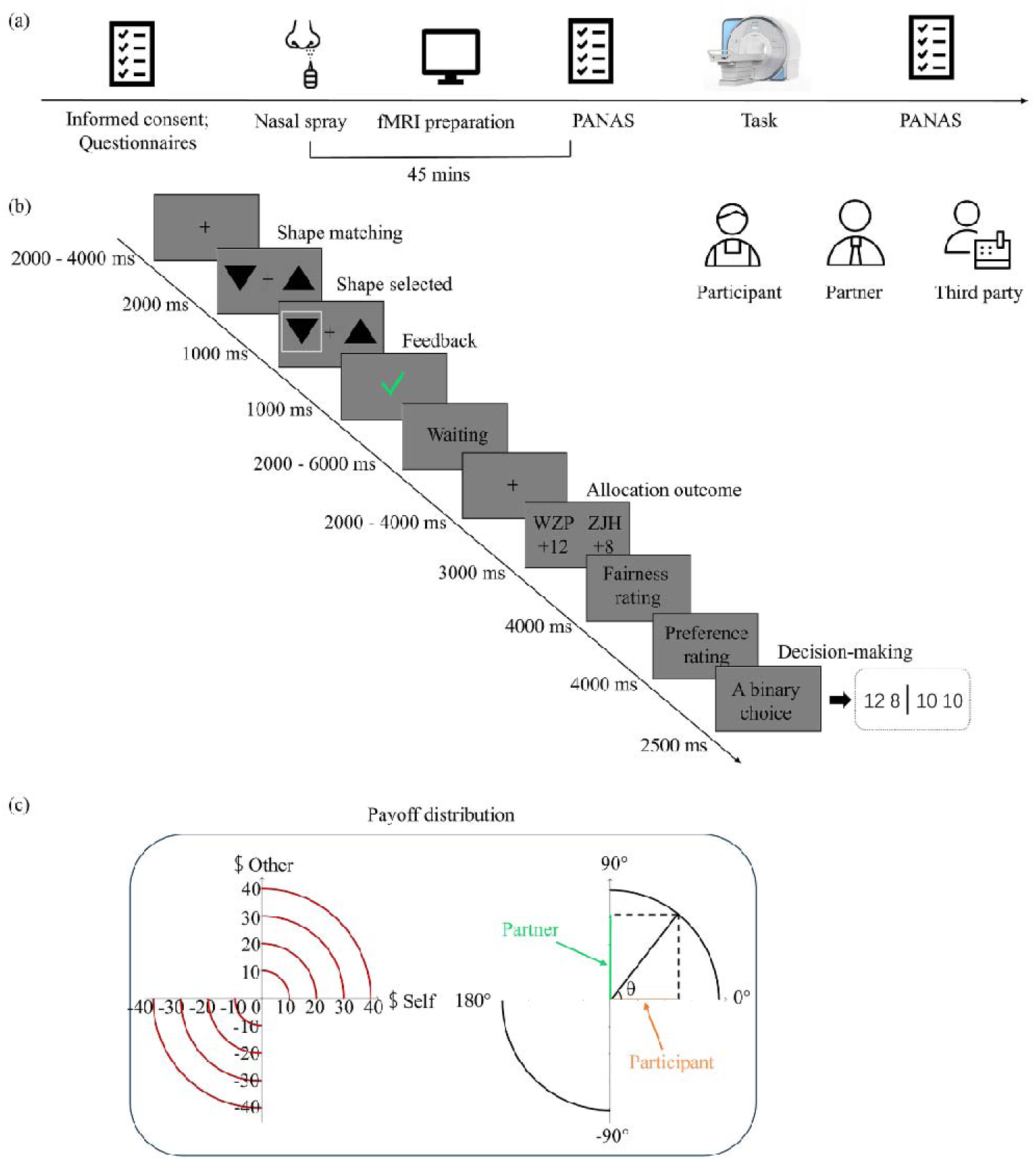
The experimental protocol and paradigm. (a) Timeline of experimental protocol. Participants first filled in their basic information, informed consent and questionnaires before administering nasal spray. The task started 45 minutes post-treatment. The Positive and Negative Affect Schedule is completed when they just arrived, 45 min post-treatment and after the fMRI scan. (b) The monetary allocation evaluation (MAE) paradigm. Participants conducted the MAE paradigm with a partner and a third party. Each trial started with a shape matching task and then a third party would allocate the gain if they matched successfully or the loss if they failed. Following presentation of the allocation outcome, participants were then required to rate their perceived levels of fairness and preference to the allocation using a 9-point Likert scale respectively. At the end of the trial, participants were asked to make a choice between the actual allocation in the current trial and a fair allocation and were informed that their choices would not change the actual allocation and their final payment. (c) A schematic illustration of how allocations in the gain and loss contexts were determined. These monetary allocations were sampled on four circles centered at the origin point (0, 0) in the Cartesian coordinate space with different circumference (radius: 10, 20, 30, or 40). To cover the gain and loss allocations, monetary pairs were only sampled from the first quadrant (0°-90°) for gain trials and form the third quadrant (180°-270°) for loss trials. Gain or loss allocations were determined by an angle θ starting from 5° with an interval of 10°. The cosine θ value of the radius represents the amount of money allocated to the participant and the sine θ value indicates the amount allocated to the partner.

## Methods and materials

### Participants and treatment

Eighty healthy participants (mean age, 20.73 ± 2.08 years) were recruited for the present double-blind, placebo-controlled, between-subject pharmacological study. Based on an a priori power analysis using the G*Power v.3.1 toolbox [59], this sample size was adequate to achieve a power > 0.80 (effect size f = 0.25, α = 0.05, ANOVA: repeated measures, within-between interaction). All participants were right-handed, had normal or corrected-to-normal vision and reported no history of psychiatric, neurological, or other medical conditions. Five participants were excluded due to their expressed disbelief of the experimental setting (3 participants) or excessive head movement (2 participants). Subsequently, 37 participants in the OT group (mean ± SD age: 20.51 ± 1.98 years) and 38 participants in the PLC group (mean ± SD age: 20.95 ± 2.17 years) were included in the final data analysis. Participants were randomly assigned to self-administered 24 IU OT or PLC following a standardized administration protocol [60] (details see Supplementary Materials-SM). Chinese versions of validated questionnaires of mood and personality traits were completed before treatment to control potential confounding effects (see SM). This study was approved by the local ethics committee at the University of Electronic Science and Technology of China and all participants provided written informed consent. All procedures were in accordance with the latest version of the Declaration of Helsinki. The study was pre-registered as a clinical trial (NCT06405737).

### Experimental paradigm

The MAE paradigm was developed based on a payoff distribution task [26]. There were three roles in the game including a third party for distributing monetary gain or loss and 2 recipients. Although participants were told that their roles were randomly distributed, they actually always acted as a recipient and the two partners were played by experimenters. At the beginning of each trial, an upward and a downward triangle were randomly presented and the participant and his “partner” were instructed to choose one from them. If they chose the same triangle, they would be rewarded (gain trials), otherwise they would be punished (loss trials). A jittered screen showing “waiting for distribution” was presented for 2-6 seconds to simulate that the third party was making allocation decisions. Unbeknown to participants, the amount of gain or loss that the third party distributed to the 2 recipients were actually pre-programmed. Allocation outcomes to the participant or his partner were indicated by their initials. Participants were then required to rate their perceived levels of fairness (1 = very unfair, 9 = very fair) and preference (1 = not at all, 9 = very preferable) to the allocation. Finally, participants were instructed to make a hypothetical choice between the proposed allocation and a fair alternative. Participants were clearly informed that their decisions would not alter the actual allocations, thereby minimizing the potential influence of participants’ decisions on their ratings, which reflects a trade-off commonly encountered in social decision-making paradigms. There were 48 gain trials and 48 loss trials, with each condition comprising 16 AI, 16 DI and 16 fair trials (details see SM).

### Data analyses

#### Behavioral analyses

Behavioral data analyses were conducted using the SPSS software (version 24.0, SPSS Inc., Chicago, IL, USA). Independent t-tests were conducted to compare group differences on questionnaire scores. For fairness and preference rating scores and the ratio of choosing unequal options, repeated-measures ANOVAs were conducted with treatment (OT vs. PLC) as between-subject factor and context (gain vs. loss) and inequity type (AI vs. DI) as within-subject factors. Greenhouse-Geisser correction was used when sphericity assumptions were violated and Bonferroni correction was applied for multiple comparisons in post-hoc analyses. Correlations between behavioral measures were examined using Pearson or Spearman coefficients depending on data distribution. Based on previous findings that subjective preferences for fairness can influence inequity-related decisions [10, 26, 50], mediation analyses were conducted to explore whether preference ratings mediated OT’s effects on fairness perception and inequity-related choices using the PROCESS macro (v4.3) (details see SM).

The Fehr-Schmidt model [5] was used to elucidate whether OT would modulate inequity aversion in different contexts. This model-based analysis was conducted based on the choice data using hierarchical Bayesian modeling with the hBayesDM package (1.2.1) [61] and Stan package (v2.32.6) [62] (see SM for model comparison and validation). All p values were based on two-sided tests.

#### fMRI data acquisition and analysis

Images were acquired on a 3.0-T GE Discovery MR scanner (General Electric Medical System, Milwaukee, WI, USA) and processed using SPM 12 (details see SM). On the first-level, different general linear models (GLMs) were designed for the outcome and decision-making phases. For the outcome phase, the GLM1 was constructed to examine neural activity in response to different types of inequity allocations and the design matrix included regressor of interest for AI and DI outcomes in the gain and loss conditions respectively. A parametric modulation analysis was conducted based on trial-by-trial fairness and preference rating scores to identify brain activity modulated by perceived levels of fairness and preference. The GLM2 was constructed to identify neural responses to different AI/DI allocations during the decision-making phase. Regressors in GLM2 were similar to GLM1 but focused on the decision-making phase in different conditions (details see SM).

On the second-level, a flexible factorial design was employed to examine the main effect of inequity type and context and the interaction effect. An independent t-test was used to test the main effect of treatment. For whole-brain level analyses, a threshold of p < 0.05 false discovery rate (FDR) corrected at peak level was set for multiple comparisons [63]. Furthermore, based on previous studies of neural substrates underpinning inequity processing [17, 26–28, 54–56], regions including the ACC, insula, dlPFC, vlPFC, medial orbitofrontal cortex (mOFC) and putamen were defined as a priori regions of interest (ROIs) to examine OT’s effects in a more sensitive way. For these a priori ROIs, the small volume correction (SVC) was applied using a threshold of p < 0.05 family-wise error (FWE) corrected on the peak level. These ROIs were anatomically defined using the third version of Automated Anatomic Labeling atlas [64, 65].

#### Representational similarity analysis

Given the advantage of representational similarity analysis (RSA) in comparing representations across multiple dimensions in higher-order space [66, 67], we examined whether OT modulated representational similarity of inequity by constructing representational dissimilarity matrices (RDMs) of behavioral ratings and aforementioned a priori ROIs associated with inequity processing for the two groups respectively. The NeuroRA toolbox (v1.1.6.12) was used to construct RDMs following a standardized procedure [66, 67]. Given that OT’s effects were found to be robust on modulating neural activity linearly encoding fairness rating scores, we therefore restricted the RSA only to fairness ratings and corresponding contrast images of each condition derived from the parametric modulation analysis of fairness ratings. For each ROI-based RDM of the two groups, the dissimilarity between one condition and another was calculated using the Pearson correlation and transformed into a measure of dissimilarity by subtracting the Pearson’s r from 1 (i.e., 1-r), resulting in a RDM that is symmetrical along its diagonal. The correlation between the two RDMs for each ROI of the two groups was calculated using the Spearman correlation [67]. A permutation test (10000 iterations) was performed to test the significance of correlations. The FDR correction was used to control for the number of ROIs used in the RSA.

## Results

### Demographics and questionnaires

Group differences of age, personality traits (see Table S1) and changes in positive and negative mood (see Table S2) were examined using independent t-tests and revealed no significant differences between the OT and PLC groups (all *ps* ≥ 0.129).

### Behavioral results

#### OT increases perceived levels of fairness and preference to AI allocations

A repeated-measures ANOVA on fairness rating scores showed significant main effects of context (F(1,73) = 42.761, *p* < 0.001, □_p_^2^ = 0.369) and inequity type (F(1,73) = 54.815, *p* < 0.001, □_p_^2^= 0.429), with participants perceiving a higher level of fairness for gain and AI allocations compared with loss and DI ones respectively. Furthermore, there was a significant treatment × inequity type interaction effect (F(1,73) = 8.680, *p* = 0.004, □_p_^2^ = 0.106; Figure 2a). Post-hoc analyses showed that while OT increased fairness rating scores to AI allocations (*p* = 0.045), it had no significant effect on DI allocations (*p* = 0.178). There were no other significant main or interaction effects (all *ps* ≥ 0.367).

**Figure 2.**
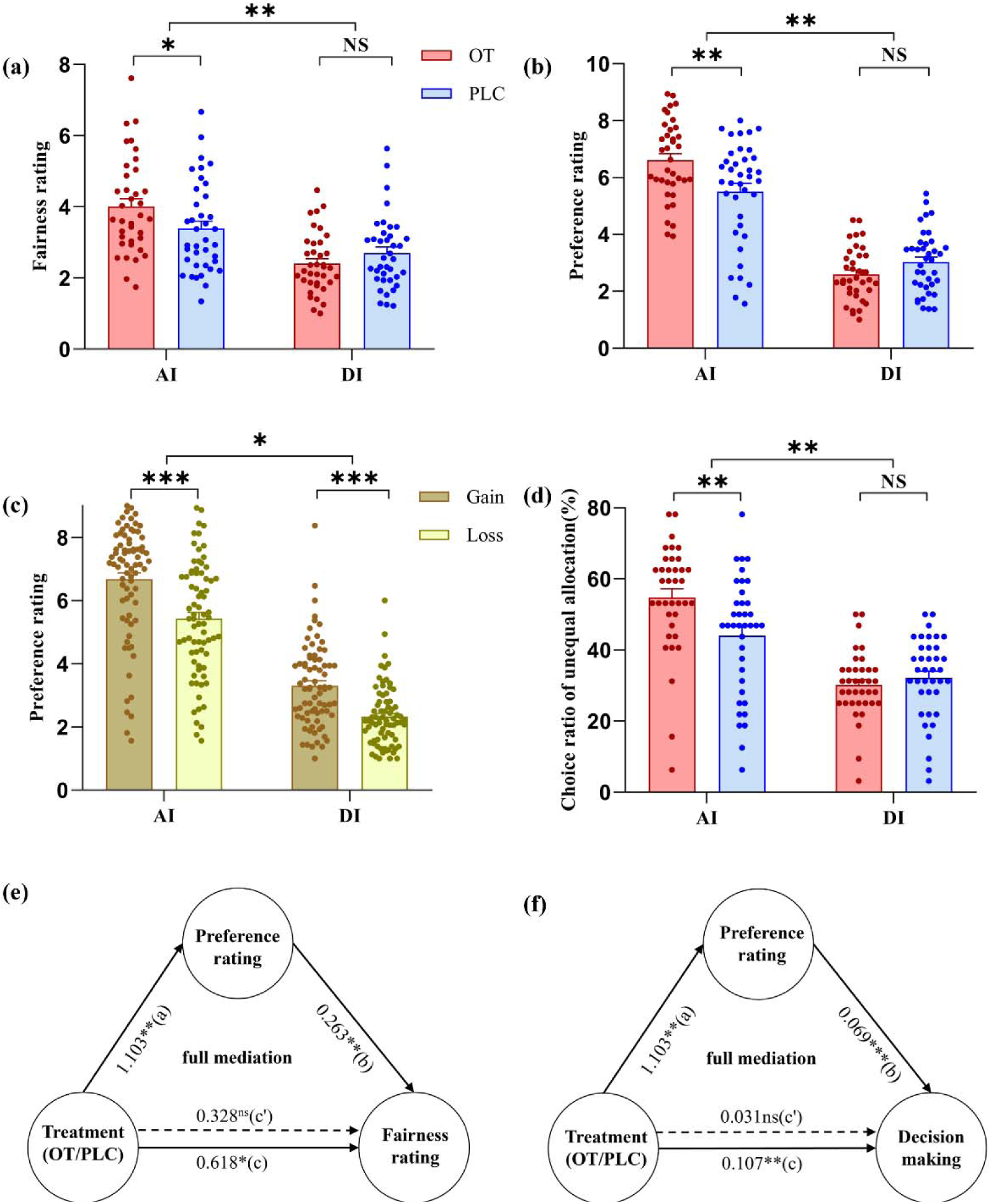
Behavioral results of ratings, choices of unequal allocations and mediation analyses. (a) Fairness ratings of oxytocin (OT) and placebo (PLC) groups in the advantageous inequity (AI) and disadvantageous inequity (DI) conditions. (b) Preference ratings of OT and PLC groups in the AI and DI conditions. (c) Preference ratings of AI and DI in the gain and loss conditions. (d) Choice ratio of unequal allocations in the AI and DI conditions. (e) Mediation analysis for treatment effects on fairness ratings via preference ratings of AI allocations. Preference rating acted as a full mediator on the association between treatment and fairness ratings. (f) Mediation analysis for treatment effects on choices of AI allocations via preference ratings. Preference rating fully mediated treatment effects on choice ratio of AI allocations (ns, not significant; * p < 0. 05, ** p < 0. 01, *** p < 0.001).

Similarly, participants showed greater preference for gain and AI allocations than to loss (main effect of context, F(1,73) = 92.553, *p* < 0.001, □_p_^2^= 0.559) and DI (main effect of inequity type, F(1,73) = 193.350, *p* < 0.001, □_p_^2^= 0.726) ones respectively. The treatment × inequity type interaction was also significant (F(1,73) = 10.861, *p* = 0.002, □_p_^2^ = 0.130; Figure 2b). Post-hoc analyses showed an enhancement effect of OT on preference for AI (*p* = 0.004) but a marginal inhibitory effect for DI allocations (*p* = 0.064). There was also a significant context × inequity type interaction (F(1,73) = 5.808, *p* = 0.018, □_p_^2^ = 0.074; Figure 2c), which was driven by a larger difference between gain and loss allocations of AI than DI (t(74) = 2.39, *p* = 0.020, Cohen’s d = 0.24). No other significant main or interaction effects were found (all *ps* ≥ 0.106).

#### OT enhances choices of AI allocations during decision-making

During decision making, participants exhibited a higher ratio of choosing unequal allocations (relative to equal allocations) in the gain than in the loss context (main effect of context, F(1,73) = 171.100, *p* < 0.001, □_p_^2^= 0.701), in the AI than in the DI condition (main effect of inequity type, F(1,73) = 67.871, *p* < 0.001, □_p_^2^= 0.482), and in the OT than in the PLC group (main effect of treatment, (F(1,73) = 3.989, *p* = 0.0495, □_p_^2^ = 0.052). Importantly, similar to ratings, the treatment × inequity type interaction was significant (F(1,73) = 8.299, *p* = 0.005, □_p_^2^ = 0.102) such that while OT increased the choice ratio of unequal allocations in the AI condition (*p* = 0.005), it had no significant effect in the DI condition (*p* = 0.413; Figure 2d). There were no other significant interaction effects (all *ps* ≥ 0.120).

#### Results of mediation and correlation analyses

Mediation analyses were conducted to explore whether preference ratings mediated OT’s effects on fairness perception (Model 1) and choices of AI allocations (Model 2). In Model 1, results showed that OT increased preference rating (path *a* = 1.103, *p* = 0.004) and fairness rating scores (path *c* = 0.618, *p* = 0.045). Preference rating scores were found to be positively associated with fairness rating scores (path *b* = 0.263, *p* = 0.005) of AI allocations. However, when preference rating was included as a mediator in the model, the direct effect of treatment on fairness ratings was not significant (path *c*′ = 0.328, *p* = 0.286). Similarly in Model 2, we found that OT also increased the choice ratio of AI allocations (path *c* = 0.107, *p* = 0.005). Preference rating scores were positively associated with the choice ratio of AI allocations (path *b* = 0.069, *p* < 0.001) and when included as a mediator, the direct effect of treatment on the choice ratio of AI allocations also became insignificant (path c′ = 0.031, *p* = 0.264). These findings suggest that the enhancement effects of OT on fairness perception (indirect effect = 0.290, SE = 0.150, 95% CI = [0.056, 0.635]; Figure 2e) and choosing AI allocations (indirect effect = 0.076, SE = 0.027, 95% CI = [0.026, 0.133]; Figure 2f) were fully mediated by participants’ preference. However, in the DI condition, there was only a significant positive correlation between fairne..ss and preference rating scores (r = 0.880, *p* < 0.001). No significant correlations were found between the choice ratio of DI allocations and either fairness (*p* = 0.431) or preference rating scores (*p* = 0.192).

#### Model-based results of OT’s effects on AI vs. DI aversion

As an independent measurement from behavioral ratings, model-based results using the Fehr-Schmidt model showed significant OT effects on decreasing AI aversion (95% HDI = [-0.67, -0.28]; Figure 3a) but increasing DI aversion (95% HDI = [0.05, 0.47]; Figure 3b). Model comparison and validation were reported in Table S3 and Figure S1. Given that hypothetical choices did not affect participants’ final payment, they may not fully reflect participants’ incentivized preference. To address this possibility, we also fitted the Fehr-Schmidt model based on preference ratings and the results replicated the current findings (see Supplemental Results and Figure S2).

**Figure 3.**
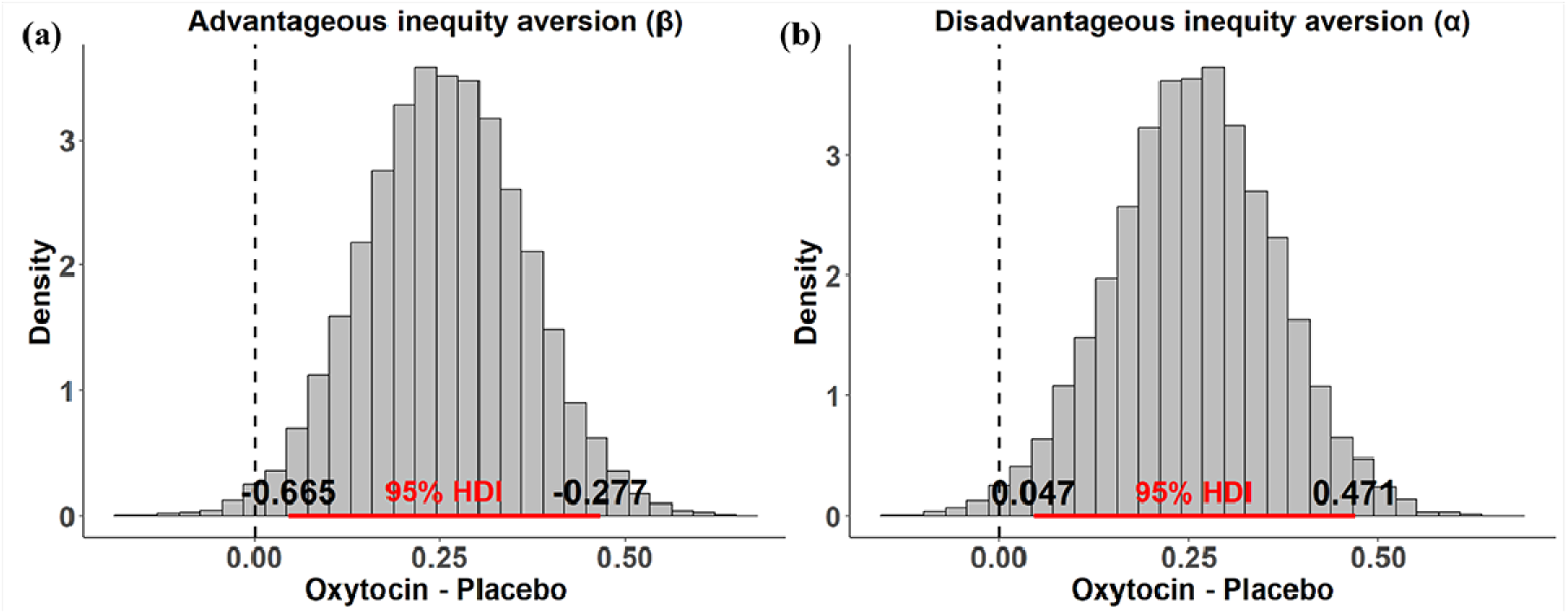
Posterior distribution and 95% highest density interval (HDI) of the group difference (oxytocin – placebo) for the inequity aversion parameter. (a) Posterior distribution and 95% HDI of the group difference for the advantageous inequity (AI) aversion parameter (β). The HDI does not include 0, suggesting that the mean parameter estimates under oxytocin are lower than under placebo, i.e., oxytocin significantly decreases AI aversion. (b) Posterior distribution and 95% HDI of the group difference for the disadvantageous inequity (DI) aversion parameter (α). The mean parameter estimates under oxytocin are greater than under placebo, i.e., oxytocin significantly increases DI aversion.

### Neural results

#### OT increases neural responses to AI allocations

To examine neural substrates encoding subjectively perceived fairness and preference, parametric modulation analyses were conducted in the outcome phase by including fairness or preference rating scores as a parametric modulator. Results at the whole-brain level showed that, similar to the behavioral pattern, OT increased neural activity associated with fairness rating scores in the bilateral dlPFC and dACC, the right dorsal aINS, vlPFC, and VS (*p*_FDR_ < 0.05; Figure 4a and Table S4) in response to AI compared with DI allocations independent of context ([OT_AI_ > PLC_AI_] > [OT_DI_ > PLC_DI_]). Further examination of each specific condition showed a similar effect of OT on neural responses to AI allocations in the loss condition (Supplementary Table S5). There were no other significant main or interaction effects at the whole brain level using the same threshold (*p*_FDR_ < 0.05).

**Figure 4.**
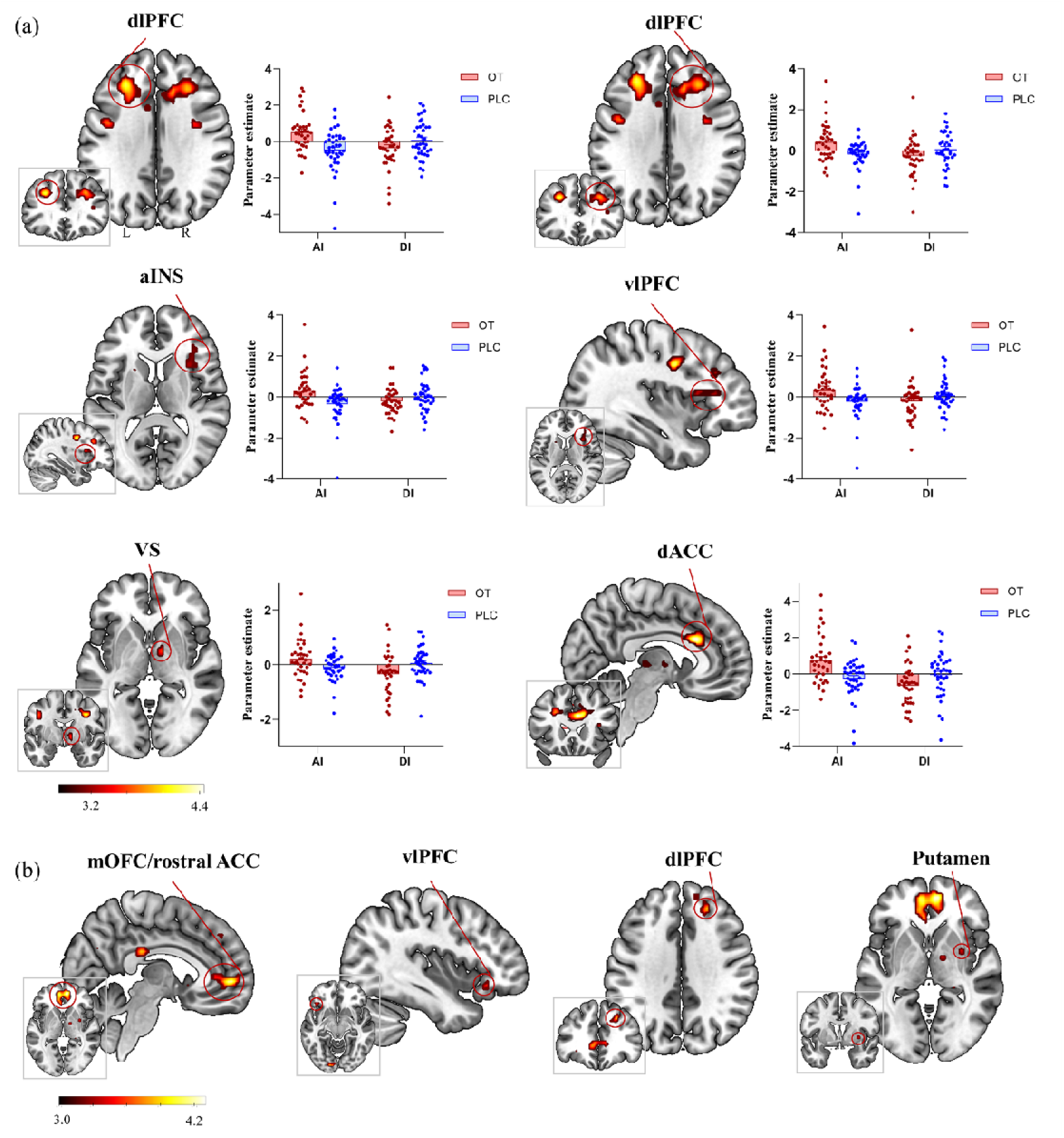
Neural results of parametric modulator analyses for fairness and preference ratings. (a) Parametric modulator analyses based on fairness rating showed that OT increased the dlPFC, left dACC and right dorsal aINS activity in response to ([OT_AI_ > PLC_AI_] > [OT_DI_ > PLC_DI_]). Statistical maps are displayed a peak-level threshold p_FDR_ < 0.05. (b) Parametric modulator analyses based on preference rating found that OT increased the activity in the right mOFC/rostral ACC, dlPFC, putamen and the left vlPFC in response to AI allocations in the gain context (OT_GainAI_ > PLC_GainAI_; p_FWE_ < 0.05, SVC). Statistical maps are displayed a peak-level uncorrected threshold P < 0.001. AI, advantageous inequity. DI, disadvantageous inequity. ACC, anterior cingulate cortex. dlPFC, dorsolateral prefrontal cortex. vlPFC, ventrolateral prefrontal cortex. aINS, anterior insula. mOFC, medial orbitofrontal cortex. VS, Ventral striatum.

For the parametric modulator of preference ratings, OT significantly increased activity in the right medial orbitofrontal cortex (mOFC)/rostral ACC (x/y/z: 6/53/-1, voxel = 212, t = 4.29, *p*_FWE_ < 0.05, SVC), dlPFC (x/y/z: 24, 38, 35, voxel = 30, t = 3.92, *p*_FWE_ < 0.05, SVC), putamen (x/y/z: 33, 2, -1, voxel = 7, t = 3.52, *p*_FWE_ < 0.05, SVC) and the left vlPFC (x/y/z: -39, 29, -10, voxel = 14, t = 3.64, *p*_FWE_ < 0.05, SVC) in response to AI gain allocations (Figure 4b). No significant main or interaction effects were found at the whole brain level using the same threshold (*p*_FDR_ < 0.05).

#### Representational similarity analysis of inequity encoding

For the RSA, to rule out the possibility that low within-group representational similarity contributed to the absence of significant correlations between the RDMs of the two groups, we first used a split-half approach (1000 iterations) to assess within-group RDM similarities at both the behavioral and neural levels (Supplementary Table S6). Only RDMs exhibiting significantly greater similarity within both the OT and PLC groups than between groups were reported (Figure 5). Results showed that the correlation between the behavioral RDMs of fairness ratings in the OT and PLC groups was not significant (r = 0.486, *p* = 0.156, permutation test with 10,000 iterations). At the neural level, we also found no significant correlations between the RDMs of the two groups for all the a priori ROIs (*p*_FDR_ ≥ 0.130).

**Figure 5.**
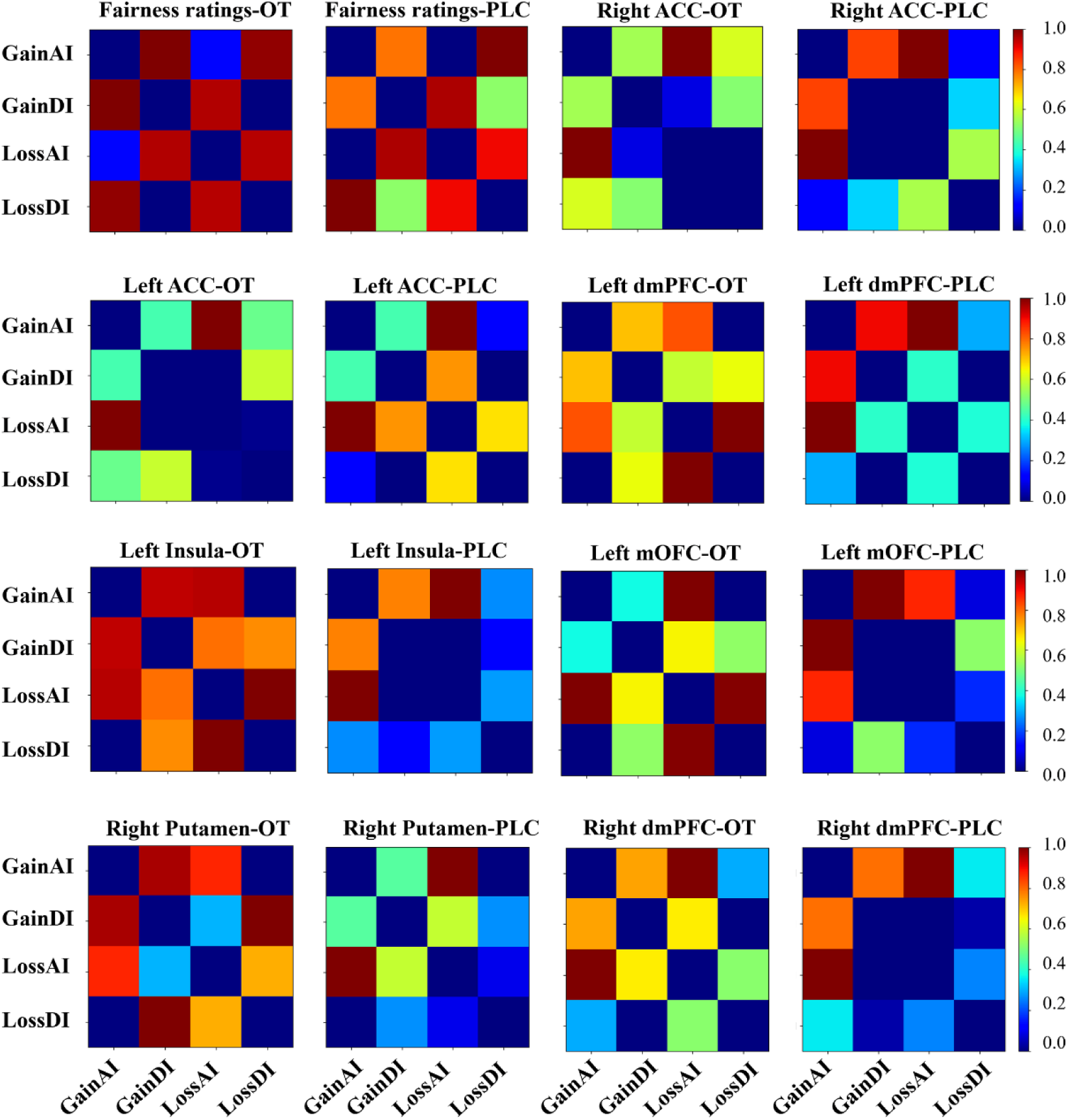
Representational dissimilarity matrices (RDMs) for fairness ratings and each ROI in both the oxytocin (OT) and placebo (PLC) groups. Regions used in the RDM analysis were the same to those in our a priori ROI analysis. Only RDMs that showed significantly higher similarity within both the OT and PLC groups than between groups after multiple comparison correction were reported. ACC, anterior cingulate cortex. dmPFC, dorsomedial prefrontal cortex. mOFC, medial orbitofrontal cortex.

#### OT modulates neural responses during decision-making

During the decision-making phase, we mainly focused on how OT modulated neural activity associated with different choices. Although we did not find significant effects associated with choosing AI or DI allocations, at the whole brain level OT was found to decrease activity in the bilateral dorsal aINS/vlPFC, dACC, the left temporal parietal junction (TPJ), putamen, and the right dlPFC, dorsomedial prefrontal cortex (dmPFC)/SMA when choosing fair allocations paired with AI allocations (i.e., participants rejected AI allocations) (*p*_FDR_ < 0.05; Figure 6a and Table S7). OT also decreased activity in the bilateral dlPFC, the left TPJ, dACC, and the right dmPFC/SMA, dorsal alNS/vlPFC when choosing fair allocations paired with DI allocations (i.e., participants rejected DI allocations) (*p*_FDR_ < 0.05; Figure 6b and Table S8). No significant OT effects were found when choosing fair allocations paired with AI compared to DI allocations (*p*_FDR_ < 0.05). Furthermore, comparison between choosing the DI allocation and its paired fair allocation showed that OT increased neural activity in the bilateral dlPFC, TPJ, precuneus/PCC, and the left vlPFC and dorsal aINS (*p*_FDR_ < 0.05; Figure 6c and Table S9). Other significant effects not related to OT were reported in the Table S10.

**Figure 6.**
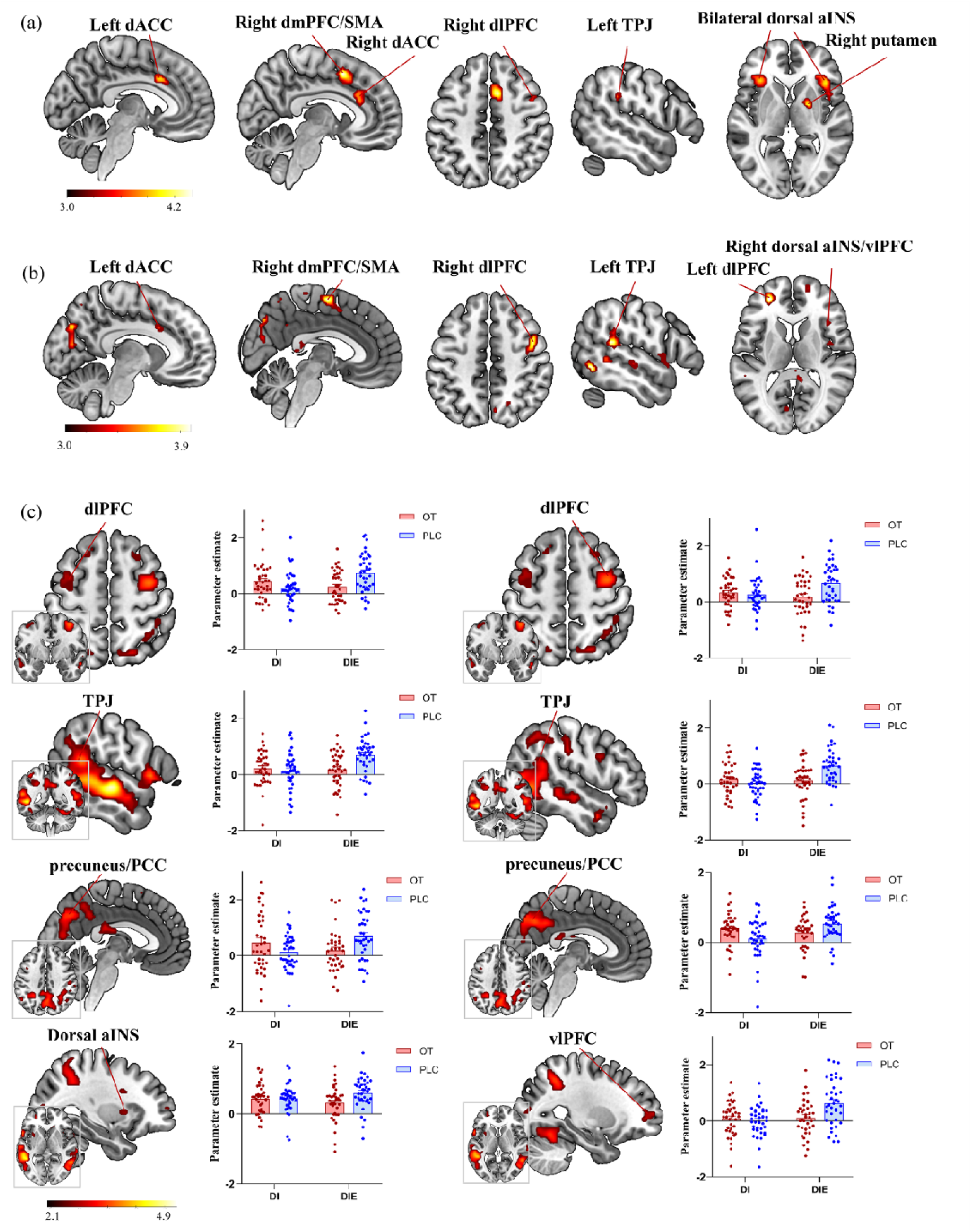
Neural effects of oxytocin (OT) associated with choices during the decision-making phase. (a) Decreased neural responses associated with choosing equal allocations paired with AI allocations (i.e., rejecting the AI allocation) in the OT compared with the placebo (PLC) group (OT_AIE_ < PLC_AIE_). (b) OT decreased neural responses associated with choosing equal allocations paired with DI allocations (i.e., rejecting the DI allocation) (OT_DIE_ < PLC_DIE_). (c) OT increased neural responses associated with choosing the DI allocation compared with choosing its paired equal allocation (OT_DI > DIE_ > PLC_DI > DIE_). Statistical maps are displayed a peak-level threshold of p_FDR_ < 0.05. AIE, equal allocations paired with AI allocations. DIE, equal allocations paired with DI allocations. AI, advantageous inequity. DI, disadvantageous inequity. ACC, anterior cingulate cortex. dlPFC, dorsolateral prefrontal cortex. dmPFC, dorsomedial prefrontal cortex. vlPFC, ventrolateral prefrontal cortex. TPJ, temporal parietal junction. SMA, Supplementary Motor Area. aINS, anterior insula. PCC, posterior cingulate.

## Discussion

Using a MAE paradigm, the present study investigated whether intranasal OT could modulate AI vs. DI perception of recipients in gain vs. loss contexts. Behavioral results showed that OT increased perceived levels of fairness and preference as well as choices in response to AI. Furthermore, both the enhancement effects of OT on fairness perception and AI choices were fully mediated by OT increasing preference for AI allocations. Computational modeling results revealed that OT decreased AI aversion but increased DI aversion. These results were paralleled by neural findings that OT increased neural activity associated with fairness ratings in brain regions engaged in social norm compliance, conflict monitoring, and inequity aversion in response to AI compared with DI allocations. These findings were further supported by multivariate based RSA at both the behavioral and neural levels.

At the behavioral level, participants perceived a higher level of fairness and preference for AI compared with DI allocations. They also chose more AI than DI allocations during decision making. These findings suggest a self-serving bias consistent with previous findings of a higher level of satisfaction with, and a lower rejection rate to, AI than DI allocations [10, 26]. In accordance with previous studies [14, 15, 26], participants also perceived gain allocations as fairer and more preferable than loss allocations. Consistently, participants chose more unequal allocations in the gain than the loss context. These findings together validated the MAE paradigm.

More importantly, OT was found to significantly increase the perceived level of fairness and preference for AI allocations independent of gain vs. loss contexts, which was maintained to the decision-making phase such that OT increased AI choices but had no significant effect on DI ones. Our mediation analyses further showed that OT’s effects on increasing the perceived fairness and AI choices were fully mediated by OT enhancing participants’ preference for AI allocations. OT therefore may modulate fairness perception and fairness-related decision-making for AI allocations via influencing individuals’ attitudes. This argument was further supported by model-based findings that OT significantly reduced AI aversion. However, for DI allocations OT was found to significantly increase DI aversion only in the model-based analysis, although its effects on behavioral ratings and choices trended in the same direction, suggesting that the model-based analysis may be more sensitive in capturing behavioral preference.

Note that these findings were inconsistent with the conventional view of OT as a prosocial hormone, particularly in light of a previous study reporting that OT reduced choices of AI but not DI allocations [48]. This inconsistency may be due to different roles participants acted. While in the present study participants acted as recipients receiving allocations determined by a third party, in Kapetaniou et al. [48] participants served as proposers in a modified DG. There is evidence showing that AI aversion, as indicated by satisfaction rates, disappears when participants play as recipients as opposed to proposers [50]. Consistently, previous studies have shown that participants are more likely to accept AI allocations when act as recipients [68, 69]. Compared with proposers, recipients experience a lower sense of agency and a weaker perceived responsibility for unequal outcomes, both of which contribute to attenuated AI aversion [50, 70, 71]. The present study therefore extends previous findings by providing evidence that OT’s effects on inequity-related behavior may depend importantly on whether individuals are evaluating allocations imposed on themselves vs. actively making allocations for others. Although OT has traditionally been established as a prosocial hormone, there is increasing evidence showing that its functional effects are highly context-dependent [30, 33, 72–74]. OT has been demonstrated to facilitate self-benefiting decisions when self-interest conflicts with altruism or in moral dilemma scenarios [44] and to increase self-serving lying [45].

These facilitatory effects of OT on AI allocations were paralleled by similar effects at the neural level. The parametric modulation analysis showed that OT increased activity associated with perceived fairness in key regions of the fairness norm enforcement circuitry, including the bilateral dlPFC and dACC and the right dorsal aINS, vlPFC and VS in response to AI relative to DI allocations [11, 75, 76]. Given that the dorsal aINS is involved in the cognitive heuristic to detect norm violations [11, 77, 78] and the dACC encodes the motivational conflict between fairness norm violations and economic self-interest [44, 79, 80], increased activity in these two regions therefore may suggest that OT sharpens the cognitive detection of norm violations and its conflict with pursuing economic self-interest of AI allocations. Given that the aINS and dACC are also core hubs of the salience network [81, 82], and in line with the salience hypothesis of OT [72], an alternative yet compatible interpretation is that increased activity in these two regions may reflect OT-enhanced salience of AI allocations or of norm conflict induced by AI allocations [33, 72]. There is ample evidence for OT modulating processing of social cues via acting on the salience network [83–87]. The dlPFC, vlPFC, and dmPFC are critically involved in emotion regulation and cognitive control [88–91]. OT-induced increases in activity within these regions may therefore reflect enhanced regulatory control that supports the pursuit of economic self-interest in AI allocations. As a support, we found OT-enhanced activity in the VS. This effect remained significant after controlling for trial-wise self-payoff, other-payoff, and difference between them (see Figure S3), suggesting that the VS may encode subjective value of AI relative to DI allocations, rather than monetary value per se. This interpretation aligns with the role of the VS in encoding not only monetary but also abstract social or even self-generated rewards [92–94]. Consistently, the VS exhibits stronger responses to AI than DI allocations [17, 54, 55, 95], even in the loss context [23, 54]. These effects of OT were more pronounced in response to AI allocations in the loss context possibly due to loss offers normally evoking stronger responses at both behavioral and neural levels [14, 15, 26]. OT has also been found to modulate activity of these lateral and medial prefrontal areas during the anticipation of monetary losses [96]. Consistent with OT-induced changes in behavioral and neural responses, using the RSA approach we obtained evidence for high within-group but low between-group representational similarity at both behavioral and neural levels. These findings together suggest that OT may primarily facilitate a bias toward AI allocations by enhancing the salience of AI-related allocations or the detection of norm violations while simultaneously strengthening regulatory processes that support the pursuit of self-beneficial outcomes.

For the parametric modulator of preference ratings, OT was found to enhance activity in response to AI gain allocations in the mOFC/rostral ACC and putamen engaged in encoding subjective positive values of rewarding stimuli [12, 26, 54–56, 58, 97] and the dlPFC and vlPFC involved in emotional regulation [88–91]. Although AI and DI allocations induced a similar trend of preference levels between the gain and loss contexts, gain AI allocations were the most preferable options (Figure 2c). It is therefore not surprising that gain AI allocations exhibit the most sensitive neural reactivity associated with preference rating scores to OT modulation. This may be achieved by OT amplifying the rewarding values of gain AI allocations through potentiating signals in the reward and emotional control systems. These neural effects were no longer significant after controlling for trial-wise self-payoff, other-payoff, and difference between them, providing further support for OT’s effects on gain AI allocations being more closely tied to the monetary values embedded in these allocations. Of note, although we observed some neural effects of OT on AI allocations selectively in the gain or loss context, we consistently observed significant interactions of treatment with inequity type but not with context at both behavioral and neural levels. OT therefore appears to modulate inequity-related processing primarily along a relative payoff imbalance dimension (i.e., AI vs. DI), rather than between gain and loss contexts, similar to the way social comparison shapes inequity perception [23, 54].

During decision-making, we focused on OT effects on neural activity associated with different choices and found that OT decreased activity in the dorsal aINS/vlPFC, dlPFC, dACC, and dmPFC engaged in cognitive control and conflict detection [11, 77, 78], and additionally in the TPJ involved in perspective taking/theory of mind [94, 98], similarly between when participants chose fair allocations paired with AI and DI allocations. Decreased activity in these regions thus may suggest reduced cognitive control and conflict processing or a decreased need to infer others’ mental state when participants decided to choose fair allocations in the OT group. OT additionally deceased activity in the putamen when choosing fair allocations paired with AI allocations, possibly indicative of it attenuating reward-seeking motivation to AI allocations [12, 93]. Furthermore, we observed an enhancement effect of OT on activation of the dlPFC, vlPFC, and dorsal aINS and additionally in the TPJ and precuneus/PCC when choosing the DI compared with its paired fair allocation. While increased dorsal aINS activation may reflect enhanced norm violation detection or DI aversion [11, 77, 78], enhanced frontal area activation may boost cognitive control to overcome such aversion for choosing DI allocations [88–91]. For the TPJ and precuneus/PCC, their stronger activation may indicate OT promoting mentalizing or inferring others’ mental states consistent with previous findings [57, 99, 100], which is also in favor of making more altruistic choices, namely choosing DI allocations.

There are several limitations in the present study. First, only males were recruited from the same cultural background and there is evidence for sex differences in approval of self-benefit behavior following OT treatment [44], thus our findings may not be extended to females. Although no culturally-dependent effects of OT have been reported to date, inferences in this regard should be made with caution. Second, given the aim of the present study, participants’ choices in the decision-making phase were hypothetical and did not affect their final payment, in order to avoid potential influences of choices on the preceding behavioral ratings (the primary outcomes). Although OT’s effects on choices were consistent with findings from behavioral ratings, computational modeling, and neural responses, inferences regarding behavioral choices and associated neural effects should take the non-incentivized nature of the choice task into consideration. Third, given the established roles of the aINS and dACC as central hubs for both salience processing and norm-violation detection, the present design did not allow us to determine whether the increased activity in these regions reflects OT-induced enhancement of the salience of AI allocations or the detection of norm violations.

In conclusion, the present neuropharmacological study provides converging evidence that OT increases AI preferences through modulation of activity in regions within the fairness-related norm enforcement circuitry. OT may act via strengthening regulatory control that supports the pursuit of self-beneficial outcomes, while simultaneously enhancing the salience of AI allocations or the detection of norm violations. We also observed preliminary evidence for OT-induced increases in DI aversion. Our findings highlight the oxytocinergic system as a key neuromodulatory candidate in shaping fairness-related behavior and provide support for both the social salience hypothesis and the hierarchical “SSS” model by offering new insights into how OT modulates human behavior through its influence on stimulus salience and subsequent higher-level processes involved in interpersonal social interactions.

## Acknowledgments

This work was supported by the National Natural Science Foundation of China (NSFC) grants (grant number: 32471139). The funders had no role in study design, data collection and analysis, decision to publish or preparation of the manuscript.

## Author contributions

S.Y. and K.M.K. designed the study. J.W., Y.Q., H.L., and H.Z. conducted the experiment and collected the data. S.Y., Y.Q., and J.W. performed the data analysis. S.Y. and J.W. wrote the manuscript draft. S.Y., Y.Z, Q.Z., and K.M.K. critically revised the manuscript draft.

## Data Availability Statement

The experimental data that support the findings of this study are available in the Open Science Framework (https://osf.io/bf738/overview?view_only=ba2431e2422849aaaf3b88c83fe9ea53).

