## Supplementary materials for "Oxytocin promotes advantageous inequity preference by modulating the fairness norm enforcement system"

**Treatment**

Participants were randomly assigned into two groups either self-administering 24 international units (IU) of OT (OT-spray, Sichuan Defeng Pharmaceutical Co. Ltd, China) or PLC (containing identical ingredients with the OT-spray other than OT, i.e., sodium chloride and glycerin) by nasal spray. They were asked to abstain from caffeine and alcohol and caffeine for 24 hours and not to have any food for 2 hours before the experiment. Following a standardized administration protocol [1], 3 puffs of OT or PLC were administered to each nostril alternately and the task began 45 minutes post-treatment. Participants were unable to guess better than chance level which treatment they had received during debriefing after the experiment (χ² = 1.07, p = 0.30).

**Questionnaires**

To control for potential confounding effects, participants completed validated Chinese versions of psychometric questionnaires on personality traits before treatment, including the Autism Spectrum Quotient [2], State-Trait Anxiety Inventory [3], Beck Depression Inventory [4, 5], Sensitivity to Punishment and Sensitivity to Reward Questionnaire [6], Behavioral Inhibition System and Behavioral Activation System Scale [7], Prosocial Tendencies Measure [8], Interpersonal Sensitivity Measure [9], the Self-report Altruism Scale [10], the Empathy Quotient [11]. For mood changes, participants completed the Positive and Negative Affect Schedule (PANAS) [12] 3 times: when they just arrived, 45 min post-treatment, and after the fMRI scan.

**Experimental Paradigm**

A monetary allocation evaluation (MAE) paradigm (Figure 1) was developed based on a payoff distribution task [13] for the present double-blind, placebo-controlled, between-subject study. Each participant came to the testing room and met two male partners. The participant was told that he would play an interactive game with the two partners in separate rooms via the local area network. There were three roles in the game including a third party for distributing monetary gain or loss and 2 recipients who were provided with an initial endowment of 20 RMB. Although participants were told that their roles were randomly distributed, they actually always acted as a recipient and the two partners were played by experimenters.

At the beginning of each trial, an upward and a downward triangle were randomly presented at the right or left side of a fixation cross and the participant and his “partner” (the other recipient) were instructed to choose one from them within 2 seconds. The selected triangle would be highlighted by a grey frame for 1 second followed by outcome feedback (matching successfully or not) for another 1 second. If they chose the same triangle, they would be rewarded (gain trials), otherwise they would be punished (loss trials). A jittered screen showing “waiting for distribution” was presented for 2-6 seconds to simulate that the third party was making allocation decisions. Unbeknown to participants, the amount of gain or loss that the third party distributed to the 2 recipients were actually pre-programmed. Allocation outcomes to the participant or his partner were indicated by their initials and were presented for 3 seconds. Participants were then required to rate their perceived levels of fairness (1 = very unfair, 9 = very fair) and preference (1 = not at all, 9 = very preferable) to the allocation using a 9-point Likert scale within 4 seconds respectively. Finally, participants were instructed to make a choice within 2.5 seconds between the actual allocation in a current trial and a fair allocation. Given that the central aim of the study was to investigate whether OT modulates how humans perceive AI vs. DI allocations and their neural correlates, participants were clearly informed that their decisions would neither alter the actual allocations nor affect their final payment, thereby minimizing the potential influence of participants’ decision consequences on their ratings. This design reflects a trade-off commonly encountered in social decision-making paradigms.

There were 48 gain trials and 48 loss trials, with each condition comprising 16 AI, 16 DI and 16 fair trials. In AI gain/loss trials, participants received greater monetary rewards/punishments than their partners (e.g., 10 vs. 3 RMB; -3 vs. -10 RMB), whereas in DI trials they received smaller monetary rewards/punishments than their partners (e.g., 3 vs. 10 RMB; -10 vs. -3 RMB). In fair trials, participants and their partners received equal monetary rewards/punishments (e.g., 10 vs. 10 RMB; -10 vs. -10 RMB; see Table S11 for details). All paired allocations were sampled on four circles centered at the origin point (0, 0) in the Cartesian coordinate space with different circumference (radius: 10, 20, 30, or 40). To cover the gain and loss allocations, monetary pairs were only sampled from the first quadrant (0°-90°) for gain trials and form the third quadrant (180°-270°) for loss trials. Gain or loss allocations were determined by an angle θ starting from 5° with an interval of 10° [14, 15]. While the cosine θ value of the radius indicates the amount of money allocated to the participant (X axis), the sine θ value indicates the amount allocated to the other recipient (Y axis) (Figure 1c). Fair trials and another four pre-programmed no-response trials were used as fillers to enhance the reality of the experiment [14, 16]. These settings led to 100 trials in total and the task lasted approximately 48 minutes. Participants performed a practice task to familiarize themselves before conducting the formal experiment. They were also informed that their final payment would be constituted by a basic participation fee and the amount of money they earned in the task. The latter was calculated as the sum of the initial endowment of 20 RMB and the allocation to the participant in a randomly selected trial [17, 18], which would be multiplied by a decimal ranging from 0.3-0.5 to avoid too much deduction (e.g., 40 RMB in the loss condition).

**Mediation analyses**

Based on previous findings that subjective preferences for fairness can influence inequity-related decisions [13, 16, 19], mediation analyses were conducted to explore whether preference ratings mediated OT’s effects on fairness perception and inequity-related choices using the PROCESS macro (v4.3). More specifically, two mediation models were constructed, with treatment as the independent variable and preference rating scores as the mediator in both models. Fairness rating scores and the choice ratio of unequal options served as the dependent variables in Model 1 and Model 2, respectively. Given that no significant group differences were found for potential confounders (e.g., age, mood and personality traits; see Tables S2 and S3), no covariates were included in these 2 models. Effects of direct and indirect paths were estimated by utilizing a bootstrapping (5000 times) method [20].

**Model-based analyses**

***The Fehr-Schmidt model***

The Fehr-Schmidt model [21] was used to elucidate whether OT would modulate inequity aversion in different contexts. This model-based analysis was conducted based on the choice data using hierarchical Bayesian modeling with the hBayesDM package (1.2.1) [22] and Stan package (v2.32.6) [23]. In this modeling, individual and group parameters (i.e., posterior distributions) are concurrently estimated in a mutually constraining manner, which makes the parameter estimation robust and reliable [22]. Four Markov Chain Monte Carlo sampling chains with 5000 iterations (including 1000 warm-up iterations) were used for parameter estimation [24]. To examine treatment effects on aversion to AI vs. DI in the gain and loss contexts, we computed the highest density interval (HDI) of the posterior distribution of the difference between group-level parameter estimates (OT vs. PLC). A HDI not including zero indicates significant group difference [22, 24, 25]. In the Fehr-Schmidt model, the subjective value of a given option is jointly determined by one's own and the other's monetary allocations according to the following formula:

*SV*(*M_self_*, *M_other_*) = *M_self_* – *α*max{*M_other_* – *M_self_*, 0} – *β*max{*M_self_* – *M_other_*, 0}, *M_self_* ≠ *M_other_*

where Mself is one's payoff and Mother is the other's payoff. Parameters α and β represent the weight assigned to DI and AI, respectively.

***Model comparison and validation***

To determine whether the Fehr–Schmidt model best fitted our data, we further compared it with another two decision-making models (Rescorla-Wagner (Delta) Model and Ideal Observer model) using the Leave-One-Out Information Criterion (LOOIC) and widely applicable information criterion (WAIC) [26]. In a totally Bayesian way, both LOOIC and WAIC estimate pointwise out-of-sample prediction accuracy, with a lower value of LOOIC or WAIC indicating higher prediction accuracy and more appropriate complexity of candidate models. LOOIC and WAIC of all candidate models were computed with the “loo” package in R [22, 26, 27]. Results indicated that the Fehr–Schmidt model was the best fitting model (see Table S4). These models were fitted using the hBayesDM package in R and all Rhat values were less than 1.1 [22].

To further validate our model’s performance, participants’ choices in each trial were simulated using posterior prediction checks of these estimated parameters in the Fehr–Schmidt model. We first plotted the stimulated data against the actual data over trials and then computed the correlation between the actual data and the stimulated data across trials. Results showed that the Fehr–Schmidt model captured the actual data well as reflected by the consistency between the actual and the stimulated data over trials and high correlations across trials (Figure S2).

**Imaging data acquisition and preprocessing**

A 3.0 Tesla GE Discovery MR750 system (General Electric Medical System, Milwaukee, WI, United States) was used to obtain T2∗-weighted echo-planar pulse sequence (TR = 2,000 ms, TE = 30 ms, slices: 43, slice thickness: 3.2 mm, field of view: 220 mm × 220 mm, matrix size: 64 × 64, flip angle: 90°). In addition, high revolution T1-weighted anatomical images were acquired obliquely with a 3D spoiled gradient echo pulse sequence (TR = 6 ms, TE = minimum, slices: 156, slice thickness: 1 mm, field of view: 256 mm × 256 mm, matrix size: 256 × 256, flip angle: 12°).

Images were processed using SPM12 software (Wellcome Department of Cognitive Neurology, London) [28]. The first five functional volumes were discarded to achieve magnet-steady images. Images were corrected for temporal slice acquisition differences and head movement using the 6-parameter rigid body algorithm. After co-registering the mean functional image and the T1 image, the T1 image was segmented to determine the parameters for normalizing the functional images to the Montreal Neurological Institute (MNI) space. The normalized images were then spatially smoothed using a Gaussian kernel (8-mm full-width at half maximum).

On the first-level, different general linear models (GLMs) were designed for the outcome and decision-making phases. For the outcome phase, the GLM1 was constructed to examine neural activity in response to different types of inequity allocations and the design matrix included regressor of interest for AI and DI outcomes in the gain and loss conditions respectively (GainAI, GainDI, LossAI, LossDI). Other regressors including the shape matching, outcome feedback, waiting screen, fairness and preference ratings, decision-making, and the six head-motion parameters were included as regressors of no interest. A parametric modulation analysis was conducted based on trial-by-trial rating scores of fairness and preference for the outcome phase to identify brain activity modulated by perceived levels of fairness and preference. The second GLM (GLM2) was constructed to identify neural responses to different AI/DI allocations during the decision-making phase. Regressors in GLM2 were similar to GLM1 but focused on the decision-making phase in different conditions.

**Model-based results of OT’s effects on AI vs. DI aversion based on preference ratings**

Given that choices in the decision-making phase did not influence participants’ payment, model-based results based on such non-incentivized choices may not reflect stable preferences. To rule out this possibility, we also fit the Fehr–Schmidt model based on preference ratings-derived utilities. To transform the continuous data of preference ratings to binary choices, we recoded them based on the median of rating scores, i.e., 5. While trials with a preference rating score higher than 5 were coded as 1 representing acceptance of the unequal option, those with a preference rating score lower than 5 were coded as 0 representing acceptance of the equal option. Trials with a rating score of 5 were deleted (an average of 4.8 trials for each participant). Results showed significant OT effects on decreasing AI aversion (95% HDI = [-0.96, -0.45]; Figure S2a) but increasing DI aversion (95% HDI = [0.37, 0.66]; Figure S2b). These results further confirmed model-based findings using choices and findings of OT effects on behavioral fairness and preference ratings based on data independent of behavioral ratings.

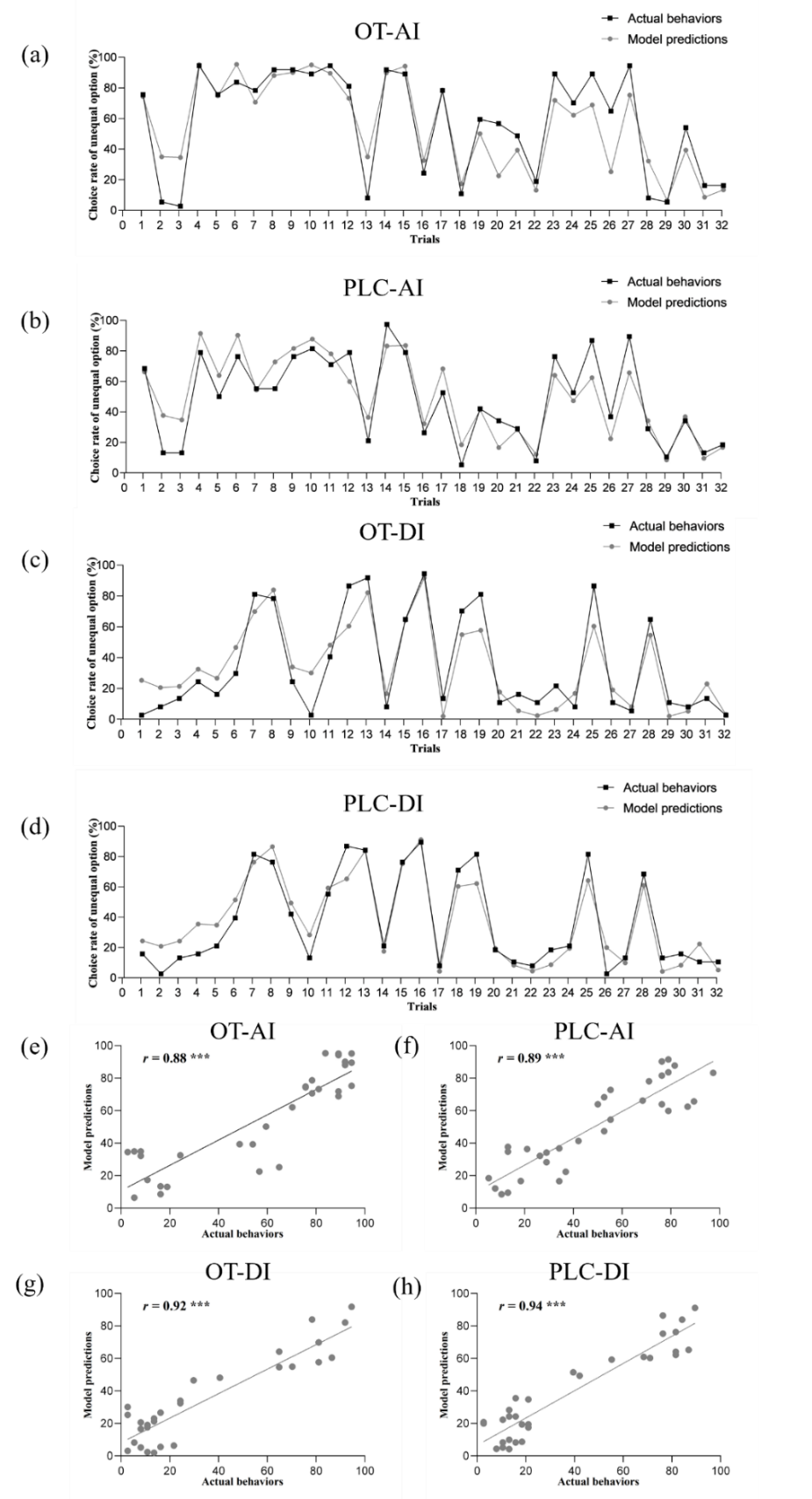

Figure S1. Results of model validation. (a-d) Consistency between actual choice rate and model-based predictions in choosing unequal options in the four conditions over trials. (e-h) Correlations between actual choice rate and model-based predictions in choosing unequal options in the four conditions across trials. ****p* < 0.001.

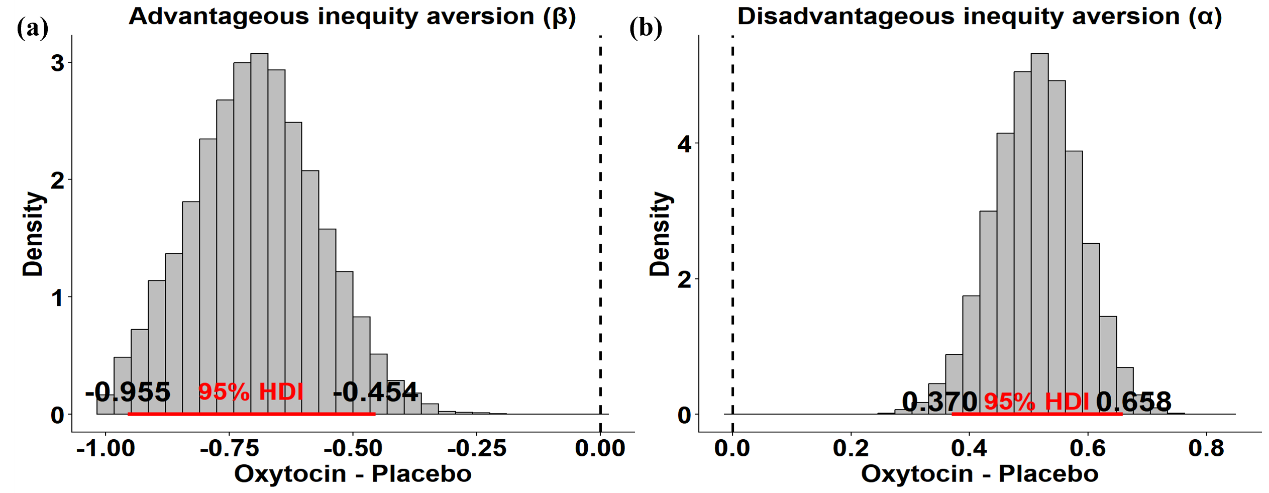

Figure S2. Posterior distribution and 95% highest density interval (HDI) of the group difference (oxytocin – placebo) for the inequity aversion parameter based on preference ratings. (a) Posterior distribution and 95% HDI of the group difference for the advantageous inequity (AI) aversion parameter (β). The HDI does not include 0, suggesting that the mean parameter estimates under oxytocin are lower than under placebo, i.e., oxytocin significantly decreases AI aversion. (b) Posterior distribution and 95% HDI of the group difference for the disadvantageous inequity (DI) aversion parameter (α). The mean parameter estimates under oxytocin are greater than under placebo, i.e., oxytocin significantly increases DI aversion. Type or paste caption here. Create a page break and paste in the Figure above the caption.

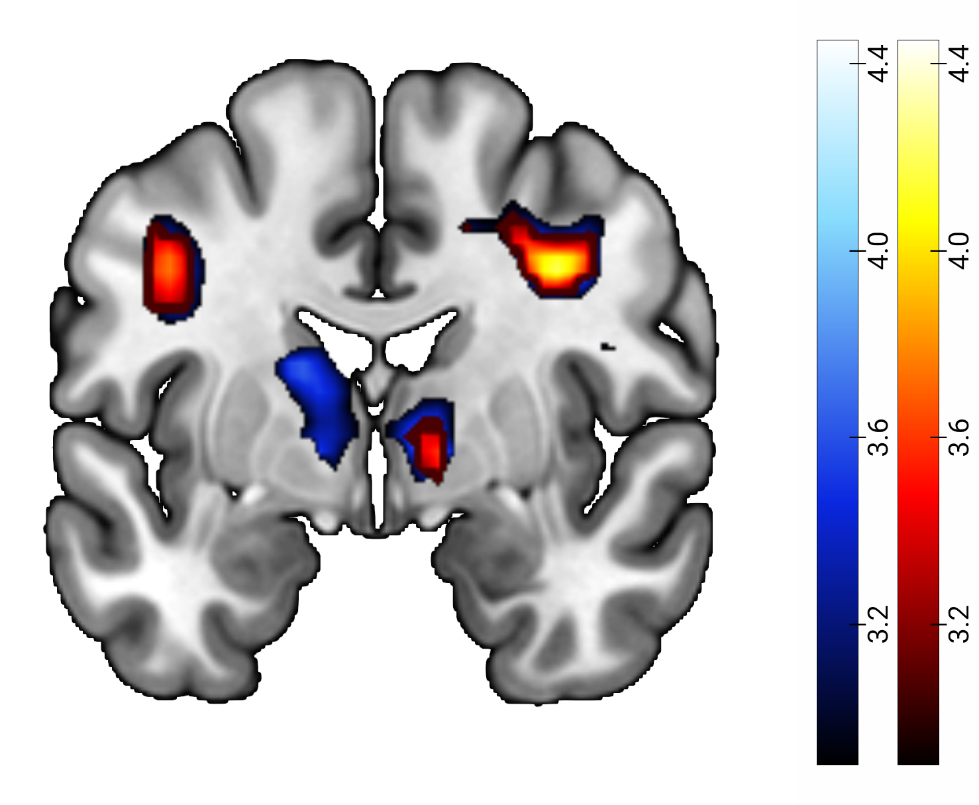

Fig. S3. Neural results of parametric modulator analyses for fairness ratings ([OT_AI_ > PLC_AI_] > [OT_DI_ > PLC_DI_]). Red represents the results of incorporating fairness ratings into the model, and blue represents the results of additionally incorporating self-payoff, other-payoff, and difference between them into the model. Significant effects in the ventral striatum were independent of whether controlling for payoffs. Statistical maps are displayed a peak-level threshold *p*_FDR_ < 0.05

Table S1. Statistics of age and questionnaire scores.

| Characteristics | OT  (M ± SD) | PLC  (M ± SD) | *t*-values | *p*-values |
| --- | --- | --- | --- | --- |
| Age | 20.51±1.98 | 20.95±2.17 | -0.904 | 0.369 |
| Autism Spectrum Quotient (ASQ) | 20.84±5.75 | 22.50±4.65 | -1.738 | 0.172 |
| Trait Anxiety Inventory (TAI) | 41.59±7.66 | 42.18±7.38 | -0.340 | 0.735 |
| State Anxiety Inventory (SAI) | 38.62±10.06 | 39.87±9.72 | -0.546 | 0.587 |
| Beck Depression Inventory (BDI-Ⅱ) | 8.24±8.09 | 8.26±5.87 | -0.012 | 0.990 |
| Sensitivity to Punishment and Sensitivity to Reward Questionnaire (SPSRQ) |  |  |  |  |
| -Sensitivity to Punishment (SP) | 9.89±3.81 | 9.76±4.25 | 0.138 | 0.891 |
| -Sensitivity to Reward (SR) | 10.14±2.98 | 9.21±3.16 | 1.302 | 0.197 |
| Behavioral Inhibition System and Behavioral Activation System Scale |  |  |  |  |
| -Behavioral Inhibition System (BAS) | 22.95±5.14 | 23.97±4.14 | -0.953 | 0.344 |
| -Behavioral Activation System (BIS) | 14.86±3.15 | 15.66±3.84 | -0.976 | 0.332 |
| Prosocial Tendencies Measure (PTM) | 69.00±13.30 | 71.87±11.17 | -1.012 | 0.315 |
| Interpersonal Sensitivity Measure (IPSM) | 62.73±12.38 | 65.16±10.28 | -0.925 | 0.358 |
| The Self-Report Altruism (SRA) | 49.84±13.45 | 47.50±8.84 | 0.887 | 0.378 |
| Interpersonal Reactivity Index-C (IRI) | 53.62±11.42 | 50.21±12.15 | 1.252 | 0.215 |
| Empathy Questionnaire (EQ) | 36.65±13.42 | 34.89±10.49 | 0.632 | 0.530 |

Table S2. Statistics of the Positive and Negative Affect Schedule (PANAS) scores.

|  | OT (M ± SD) | PLC (M ± SD) | *t*-values | *p*-values |
| --- | --- | --- | --- | --- |
| PANAS (pre-treatment) |  |  |  |  |
| -Positive | 26.65±7.32 | 24.89±6.46 | 1.101 | 0.275 |
| -Negative | 14.38±5.09 | 14.26±5.38 | 0.095 | 0.924 |
| PANAS (post-treatment) |  |  |  |  |
| -Positive | 26.70±7.49 | 24.10±7.19 | 1.532 | 0.130 |
| -Negative | 13.59±5.39 | 13.68±5.15 | -0.074 | 0.942 |
| PANAS (post-scanning) |  |  |  |  |
| -Positive | 24.08±7.69 | 21.42±7.32 | -0.312 | 0.756 |
| -Negative | 13.41±4.83 | 13.76±5.11 | 1.535 | 0.129 |

Table S3. Model comparison.

| Model | LOOIC | WAIC |
| --- | --- | --- |
| Ideal Observer model | 1408.28 | 1397.54 |
| Rescorla-Wagner (Delta) model | 1346.56 | 1338.62 |
| Fehr-Schmidt inequity aversion model | 1117.40 | 1101.10 |

Table S4. Brain regions showing increased activity induced by oxytocin in response to advantageous compared to disadvantageous inequity.

| Brain region | voxels | Peak-t value | x | y | z |
| --- | --- | --- | --- | --- | --- |
| R. Dorsal anterior cingulate | 495 | 4.49 | 6 | 23 | 26 |
| Dorsal anterior cingulate |  | 4.25 | -9 | 11 | 26 |
| Dorsolateral prefrontal cortex |  | 4.23 | -27 | 32 | 32 |
| Dorsolateral prefrontal cortex |  | 3.96 | 27 | 29 | 32 |
| R. Dorsolateral prefrontal cortex | 124 | 4.31 | 45 | 5 | 44 |
| Dorsolateral prefrontal cortex |  | 4.25 | 36 | -1 | 38 |
| L. Dorsolateral prefrontal cortex | 60 | 4.04 | -42 | -1 | 38 |
| L. Postcentral gyrus | 144 | 3.91 | -42 | -25 | 56 |
| Precentral gyrus |  | 3.89 | -30 | -28 | 62 |
| R. Inferior frontal gyrus | 17 | 3.62 | 48 | -1 | 20 |
| R. Ventral striatum | 32 | 3.61 | 12 | -4 | -1 |
| L. Dorsomedial prefrontal cortex | 20 | 3.55 | -18 | 2 | 56 |
| R. Dorsal anterior insula | 39 | 3.46 | 33 | 14 | 11 |
| Ventrolateral prefrontal cortex |  | 3.40 | 36 | 26 | 11 |
| L. Inferior occipital gyrus | 13 | 3.40 | -36 | -85 | -7 |
| R. Thalamus | 20 | 3.40 | 12 | -28 | 2 |
| Thalamus |  | 3.19 | 6 | -22 | 2 |
| L. Precuneus | 13 | 3.39 | -18 | -61 | 41 |
| L. Thalamus | 25 | 3.36 | -6 | -19 | 2 |

All with a *p*_FDR_ < 0.05 threshold and cluster > 10 voxels. L indicates left; R indicates right.

Table S5. Significant oxytocin’s effects in response to AI allocations in the loss condition in the parametric modulator analyses based on fairness rating.

| Brain region | voxels | Peak-t value | x | y | z |
| --- | --- | --- | --- | --- | --- |
| L. Dorsolateral prefrontal cortex | 146 | 4.73 | -27 | -19 | 50 |
| Dorsolateral prefrontal cortex |  | 4.15 | -27 | 29 | 32 |
| R. Dorsolateral prefrontal cortex | 126 | 3.89 | 42 | 2 | 47 |
| L. Posterior insula | 16 | 3.89 | -39 | -40 | 20 |
| L. Precentral gyrus | 35 | 3.73 | -51 | -13 | 11 |
| R. Dorsolateral prefrontal cortex | 15 | 3.40 | 36 | 50 | 14 |
| L. Middle cingulate gyrus | 16 | 3.32 | -9 | -25 | 41 |
| R. Dorsomedial prefrontal cortex | 99 | 3.74 | 3 | 32 | 41 |
| R. Anterior cingulate | 12 | 3.55 | 11 | 21 | 28 |

All with a *p*_FDR_ < 0.05 threshold and cluster > 10 voxels. L indicates left; R indicates right.

Table S6. Within-group and between-group similarities for the RDMs in both groups.

| Measures | Between groups | | Within the OT group | | | | Within the PLC group | | | |
| --- | --- | --- | --- | --- | --- | --- | --- | --- | --- | --- |
|  | r | p | r | p | z | p | PLC | p | z | p |
| **Behavior** | 0.49 | 0.156 | 0.95 | <0.001 | 6.227 | **<0.001** | 0.81 | <0.001 | 2.868 | **0.004** |
| **ACC_L** | -0.14 | 0.918 | 0.70 | 0.012 | 4.845 | **<0.001** | 0.87 | 0.002 | 7.153 | **<0.001** |
| **ACC_R** | 0.43 | 0.762 | 0.83 | 0.003 | 3.500 | **<0.001** | 0.83 | 0.003 | 3.534 | **<0.001** |
| dlPFC_L | 0.94 | 0.130 | 0.96 | <0.001 | 0.999 | 0.342 | 0.97 | <0.001 | 1.719 | 0.120 |
| dlPFC_R | 0.31 | 0.762 | 0.60 | 0.022 | 1.791 | 0.093 | 0.89 | 0.001 | 5.345 | <0.001 |
| **dmPFC_L** | 0.43 | 0.762 | 0.89 | 0.001 | 4.623 | **<0.001** | 0.83 | 0.003 | 3.534 | **<0.001** |
| **dmPFC_R** | 0.60 | 0.582 | 0.94 | 0.001 | 5.021 | **<0.001** | 0.96 | <0.001 | 6.080 | **<0.001** |
| **Insula_L** | 0.60 | 0.582 | 0.83 | 0.003 | 2.379 | **0.030** | 0.89 | 0.001 | 3.537 | **<0.001** |
| Insula_R | 0.03 | 0.957 | 0.39 | 0.045 | 1.835 | 0.093 | 0.96 | <0.001 | 9.298 | <0.001 |
| Putamen_L | 0.71 | 0.518 | 0.83 | 0.003 | 1.446 | 0.173 | 0.80 | 0.003 | 1.026 | 0.356 |
| **Putamen_R** | 0.31 | 0.762 | 0.96 | <0.001 | 7.811 | **<0.001** | 0.83 | 0.003 | 4.210 | **<0.001** |
| **mOFC_L** | 0.09 | 0.939 | 0.87 | 0.002 | 5.973 | **<0.001** | 0.94 | <0.001 | 7.997 | **<0.001** |
| mOFC_R | 0.14 | 0.918 | 0.89 | 0.001 | 6.156 | <0.001 | 0.42 | 0.042 | 1.489 | 0.174 |
| vlPFC_L | 0.31 | 0.762 | 0.62 | 0.018 | 1.944 | 0.081 | 0.39 | 0.046 | 0.443 | 0.658 |
| vlPFC_R | 0.77 | 0.504 | 0.83 | 0.003 | 0.806 | 0.420 | 0.83 | 0.003 | 0.814 | 0.447 |

L indicates left; R indicates right. OT: oxytocin. PLC: placebo. Fisher z-transformation tests were used to examine correlation differences. Measures in bold indicates those showing significantly greater similarity within both the OT and PLC groups than between groups.

Table S7. Oxytocin decreased brain activations when choosing equal allocations paired with AI allocations.

| Brain region | voxels | Peak-t value | x | y | z |
| --- | --- | --- | --- | --- | --- |
| R. Dorsomedial prefrontal cortex | 204 | 4.44 | 6 | 11 | 50 |
| Dorsal anterior cingulate |  | 4.01 | -6 | 14 | 35 |
| Dorsal anterior cingulate |  | 3.83 | 6 | 26 | 29 |
| R. Putamen | 37 | 4.17 | 21 | -1 | 2 |
| L. Supramarginal gyrus | 32 | 4.15 | -63 | -31 | 35 |
| Temporoparietal junction |  | 3.89 | -54 | -37 | 29 |
| R. Ventrolateral prefrontal cortex | 188 | 4.09 | 39 | 23 | 5 |
| Ventrolateral prefrontal cortex |  | 3.85 | 36 | 32 | -1 |
| Ventrolateral prefrontal cortex |  | 3.84 | 48 | 17 | 11 |
| L. Anterior insula | 104 | 4.00 | -33 | 26 | 2 |
| Anterior insula |  | 3.91 | -30 | 14 | -7 |
| Ventrolateral prefrontal cortex |  | 3.40 | -36 | 41 | -7 |
| R. Dorsolateral prefrontal cortex | 10 | 3.68 | 45 | 8 | 47 |

All with a *p*_FDR_ < 0.05 threshold and cluster > 10 voxels. L indicates left; R indicates right.

Table S8. Oxytocin decreased brain activations when choosing equal allocations paired with DI allocations.

| Brain region | voxels | Peak-t value | x | y | z |
| --- | --- | --- | --- | --- | --- |
| L. Dorsolateral prefrontal cortex | 97 | 4.79 | -27 | 47 | 14 |
| Dorsolateral prefrontal cortex |  | 3.46 | -27 | 50 | 2 |
| L. Middle occipital gyrus | 137 | 4.67 | -51 | -64 | -10 |
| Middle temporal gyrus |  | 3.70 | -48 | -46 | -1 |
| Middle temporal gyrus |  | 3.36 | -57 | -49 | 2 |
| L. Superior temporal gyrus | 37 | 4.30 | -57 | -22 | -4 |
| R. Superior occipital gyrus | 335 | 4.23 | 21 | -73 | 29 |
| Precuneus |  | 4.02 | -3 | -76 | 38 |
| Precuneus |  | 3.60 | 6 | -70 | 50 |
| L. Superior temporal gyrus | 70 | 4.11 | -57 | -43 | 17 |
| Supramarginal gyrus |  | 3.45 | -51 | -40 | 26 |
| Posterior insula |  | 3.35 | -45 | -34 | 20 |
| Temporoparietal junction |  | 3.26 | -54 | -40 | 23 |
| R. Dorsomedial prefrontal cortex | 83 | 4.01 | 3 | -7 | 62 |
| R. Dorsolateral prefrontal cortex | 112 | 3.99 | 42 | -4 | 56 |
| Dorsolateral prefrontal cortex |  | 3.90 | 45 | 2 | 47 |
| Dorsolateral prefrontal cortex |  | 3.49 | 30 | -4 | 59 |
| L. Ventrolateral prefrontal cortex | 40 | 3.71 | -60 | 8 | 5 |
| L. Superior parietal lobe | 13 | 3.70 | -18 | -67 | 53 |
| R. Fusiform gyrus | 46 | 3.70 | 24 | -58 | -13 |
| R. Superior temporal gyrus | 15 | 3.70 | 54 | 14 | -10 |
| R. Dorsolateral prefrontal cortex | 23 | 3.60 | 18 | 59 | 14 |
| L. Parahippocampa gyrus | 36 | 3.59 | -21 | -58 | -10 |
| R. Postcentral gyrus | 21 | 3.57 | 42 | -37 | 59 |
| R. Fusiform gyrus | 16 | 3.56 | 39 | -43 | -19 |
| L. Dorsal anterior cingulate | 10 | 3.53 | -9 | 14 | 32 |
| R. Middle cingulate gyrus | 10 | 3.52 | 6 | -49 | 35 |
| R. Precentral gyrus | 15 | 3.52 | 63 | 5 | 26 |
| L. Superior parietal lobe | 22 | 3.44 | -36 | -46 | 62 |
| R. Insula | 12 | 3.44 | 45 | -4 | 14 |
| R. Ventrolateral prefrontal cortex | 11 | 3.32 | 39 | 29 | 5 |
| Ventrolateral prefrontal cortex |  | 3.19 | 39 | 20 | 11 |

All with a *p*_FDR_ < 0.05 threshold and cluster > 10 voxels. L indicates left; R indicates right.

Table S9. Oxytocin increased brain activations when choosing DI allocations compared with choosing its paired equal allocations.

| Brain region | voxels | Peak-t value | x | y | z |
| --- | --- | --- | --- | --- | --- |
| L. Superior temporal gyrus | 1659 | 5.24 | -60 | -19 | -4 |
| Middle temporal gyrus |  | 4.54 | -48 | -46 | 2 |
| Middle temporal gyrus |  | 4.43 | -54 | -34 | -1 |
| Temporoparietal junction |  | 3.85 | -57 | -46 | 17 |
| R. Middle temporal gyrus | 1246 | 4.26 | 60 | -13 | -16 |
| Middle temporal gyrus |  | 4.10 | 54 | 2 | -28 |
| Inferior temporal gyrus |  | 4.01 | 57 | -58 | -7 |
| Temporoparietal junction |  | 3.81 | 54 | -52 | 8 |
| Temporoparietal junction |  | 3.29 | 49 | -47 | 18 |
| L. Dorsolateral prefrontal cortex | 43 | 4.06 | -15 | 14 | 62 |
| Dorsolateral prefrontal cortex |  | 2.94 | -21 | 20 | 56 |
| Dorsolateral prefrontal cortex |  | 2.66 | -21 | -1 | 65 |
| R. Inferior parietal lobule | 1921 | 3.96 | 9 | -34 | 20 |
| Precuneus |  | 3.61 | 6 | -52 | 35 |
| Precuneus |  | 3.59 | 3 | -61 | 38 |
| Precuneus |  | 3.44 | -3 | -68 | 38 |
| R. Dorsolateral prefrontal cortex | 178 | 3.78 | 36 | -4 | 56 |
| L. Precentral gyrus | 101 | 3.73 | -45 | 5 | 50 |
| Precentral gyrus |  | 2.76 | -39 | 8 | 38 |
| Precentral gyrus |  | 2.61 | -33 | -10 | 62 |
| L. Superior temporal gyrus | 22 | 3.65 | -48 | 11 | -28 |
| R. Dorsolateral prefrontal cortex | 12 | 3.35 | 21 | 20 | 59 |
| L. Dorsolateral prefrontal cortex | 33 | 3.02 | -27 | 50 | 2 |
| Dorsolateral prefrontal cortex |  | 2.95 | -27 | 59 | 5 |
| Ventrolateral prefrontal cortex |  | 2.89 | -27 | 53 | 5 |
| R. Ventrolateral prefrontal cortex | 36 | 2.97 | 54 | 8 | 26 |
| R. Dorsolateral prefrontal cortex | 20 | 2.95 | 39 | 26 | 44 |
| L. Dorsal anterior insula | 11 | 2.73 | -30 | 8 | 2 |

All with a *p*_FDR_ < 0.05 threshold and cluster > 10 voxels. L indicates left; R indicates right.

Table S10. Brain activations when choosing equal allocations compared with choosing unequal allocations.

| Brain region | voxels | Peak-t value | x | y | z |
| --- | --- | --- | --- | --- | --- |
| L. Ventrolateral prefrontal cortex | 4957 | 6.83 | -45 | 23 | -4 |
| Dorsomedial prefrontal cortex |  | 6.31 | 0 | 47 | 32 |
| Supplementary motor area |  | 6.16 | -9 | 17 | 59 |
| R. Middle temporal gyrus | 229 | 4.98 | 48 | 5 | -34 |
| Middle temporal gyrus |  | 3.66 | 51 | -13 | -13 |
| Middle temporal gyrus |  | 3.55 | 48 | -34 | -1 |
| R. Angular | 357 | 4.76 | 57 | -58 | 26 |
| Middle temporal gyrus |  | 3.22 | 33 | -64 | 26 |
| Precuneus |  | 2.80 | 30 | -64 | 41 |
| R. Inferior occipital gyrus | 241 | 4.33 | 39 | -85 | -7 |
| Middle occipital gyrus |  | 3.27 | 36 | -85 | 14 |
| L. Caudate | 54 | 3.36 | -12 | 2 | 14 |
| L. Precuneus | 126 | 3.32 | 0 | -61 | 35 |
| R. Precuneus |  | 3.29 | 3 | -55 | 23 |
| L. Superior parietal lobule | 111 | 3.31 | -24 | -58 | 44 |
| Superior parietal lobule |  | 3.27 | -27 | -70 | 53 |
| R. Hippocampus | 11 | 2.97 | 21 | -28 | -7 |
| L. Middle cingulate gyrus | 11 | 2.80 | 0 | -22 | 35 |

All with a *p*_FDR_ < 0.05 threshold and cluster > 10 voxels. L indicates left; R indicates right.

Table S11. Allocations used in the monetary allocation evaluation paradigm.

| Self | Other | Number of trials | Self vs. Other | Outcome | Inequity type |
| --- | --- | --- | --- | --- | --- |
| 8 | 6 | 1 | 2 | Win | Advantageous |
| 16 | 11 | 1 | 5 | Win | Advantageous |
| 9 | 4 | 1 | 5 | Win | Advantageous |
| 10 | 3 | 1 | 7 | Win | Advantageous |
| 25 | 17 | 1 | 8 | Win | Advantageous |
| 10 | 1 | 1 | 9 | Win | Advantageous |
| 18 | 8 | 1 | 10 | Win | Advantageous |
| 33 | 23 | 1 | 10 | Win | Advantageous |
| 20 | 2 | 1 | 12 | Win | Advantageous |
| 19 | 5 | 1 | 14 | Win | Advantageous |
| 27 | 13 | 1 | 14 | Win | Advantageous |
| 36 | 17 | 1 | 19 | Win | Advantageous |
| 29 | 8 | 1 | 21 | Win | Advantageous |
| 30 | 3 | 1 | 27 | Win | Advantageous |
| 39 | 10 | 1 | 29 | Win | Advantageous |
| 40 | 3 | 1 | 37 | Win | Advantageous |
| 7 | 7 | 1 | 0 | Win | Equal |
| 14 | 14 | 1 | 0 | Win | Equal |
| 21 | 21 | 1 | 0 | Win | Equal |
| 28 | 28 | 1 | 0 | Win | Equal |
| 8 | 8 | 1 | 0 | Win | Equal |
| 10 | 10 | 1 | 0 | Win | Equal |
| 11 | 11 | 1 | 0 | Win | Equal |
| 13 | 13 | 1 | 0 | Win | Equal |
| 15 | 15 | 1 | 0 | Win | Equal |
| 17 | 17 | 1 | 0 | Win | Equal |
| 18 | 18 | 1 | 0 | Win | Equal |
| 20 | 20 | 1 | 0 | Win | Equal |
| 22 | 22 | 1 | 0 | Win | Equal |
| 23 | 23 | 1 | 0 | Win | Equal |
| 25 | 25 | 1 | 0 | Win | Equal |
| 26 | 26 | 1 | 0 | Win | Equal |
| 6 | 8 | 1 | -2 | Win | Disadvantageous |
| 4 | 9 | 1 | -5 | Win | Disadvantageous |
| 11 | 16 | 1 | -5 | Win | Disadvantageous |
| 3 | 10 | 1 | -7 | Win | Disadvantageous |
| 17 | 25 | 1 | -7 | Win | Disadvantageous |
| 1 | 10 | 1 | -9 | Win | Disadvantageous |
| 8 | 18 | 1 | -10 | Win | Disadvantageous |
| 23 | 33 | 1 | -10 | Win | Disadvantageous |
| 5 | 19 | 1 | -14 | Win | Disadvantageous |
| 13 | 27 | 1 | -15 | Win | Disadvantageous |
| 2 | 20 | 1 | -18 | Win | Disadvantageous |
| 17 | 36 | 1 | -19 | Win | Disadvantageous |
| 8 | 29 | 1 | -21 | Win | Disadvantageous |
| 3 | 30 | 1 | -27 | Win | Disadvantageous |
| 10 | 39 | 1 | -28 | Win | Disadvantageous |
| 3 | 40 | 1 | -36 | Win | Disadvantageous |
| -6 | -8 | 1 | 2 | Loss | Advantageous |
| -4 | -9 | 1 | 5 | Loss | Advantageous |
| -11 | -16 | 1 | 5 | Loss | Advantageous |
| -3 | -10 | 1 | 7 | Loss | Advantageous |
| -17 | -25 | 1 | 7 | Loss | Advantageous |
| -1 | -10 | 1 | 9 | Loss | Advantageous |
| -8 | -18 | 1 | 10 | Loss | Advantageous |
| -23 | -33 | 1 | 10 | Loss | Advantageous |
| -5 | -19 | 1 | 14 | Loss | Advantageous |
| -13 | -27 | 1 | 15 | Loss | Advantageous |
| -2 | -20 | 1 | 18 | Loss | Advantageous |
| -17 | -36 | 1 | 19 | Loss | Advantageous |
| -8 | -29 | 1 | 21 | Loss | Advantageous |
| -3 | -30 | 1 | 27 | Loss | Advantageous |
| -10 | -39 | 1 | 28 | Loss | Advantageous |
| -3 | -40 | 1 | 36 | Loss | Advantageous |
| -7 | -7 | 1 | 0 | Loss | Equal |
| -14 | -14 | 1 | 0 | Loss | Equal |
| -21 | -21 | 1 | 0 | Loss | Equal |
| -28 | -28 | 1 | 0 | Loss | Equal |
| -8 | -8 | 1 | 0 | Loss | Equal |
| -10 | -10 | 1 | 0 | Loss | Equal |
| -11 | -11 | 1 | 0 | Loss | Equal |
| -13 | -13 | 1 | 0 | Loss | Equal |
| -15 | -15 | 1 | 0 | Loss | Equal |
| -17 | -17 | 1 | 0 | Loss | Equal |
| -18 | -18 | 1 | 0 | Loss | Equal |
| -20 | -20 | 1 | 0 | Loss | Equal |
| -22 | -22 | 1 | 0 | Loss | Equal |
| -23 | -23 | 1 | 0 | Loss | Equal |
| -25 | -25 | 1 | 0 | Loss | Equal |
| -26 | -26 | 1 | 0 | Loss | Equal |
| -8 | -6 | 1 | -2 | Loss | Disadvantageous |
| -9 | -4 | 1 | -5 | Loss | Disadvantageous |
| -16 | -11 | 1 | -5 | Loss | Disadvantageous |
| -10 | -3 | 1 | -7 | Loss | Disadvantageous |
| -25 | -17 | 1 | -8 | Loss | Disadvantageous |
| -10 | -1 | 1 | -9 | Loss | Disadvantageous |
| -18 | -8 | 1 | -10 | Loss | Disadvantageous |
| -33 | -23 | 1 | -10 | Loss | Disadvantageous |
| -19 | -5 | 1 | -14 | Loss | Disadvantageous |
| -27 | -13 | 1 | -14 | Loss | Disadvantageous |
| -20 | -2 | 1 | -18 | Loss | Disadvantageous |
| -36 | -17 | 1 | -19 | Loss | Disadvantageous |
| -29 | -8 | 1 | -21 | Loss | Disadvantageous |
| -30 | -3 | 1 | -27 | Loss | Disadvantageous |
| -39 | -10 | 1 | -29 | Loss | Disadvantageous |
| -40 | -3 | 1 | -37 | Loss | Disadvantageous |
